# Hydrodynamics of spike proteins dictate a transport-affinity competition for SARS-CoV-2 and other enveloped viruses

**DOI:** 10.1101/2022.01.03.474721

**Authors:** Nicolas Moreno, Daniela Moreno-Chaparro, Florencio Balboa Usabiaga, Marco Ellero

## Abstract

Many viruses, such as SARS-CoV-2 or Influenza, possess spike-decorated envelopes. Depending on the virus type, a large variability is present in spikes number, morphology and reactivity, which remains generally unexplained. Since viruses’ transmissibility depend on features beyond their genetic sequence, new tools are required to discern the effects of spikes functionality, interaction, and morphology. Here, we postulate the relevance of hydrodynamic interactions in the viral infectivity of enveloped viruses and propose micro-rheological characterization as a platform for viruses differentiation. To understand how the spikes affect virion mobility and infectivity, we investigate the diffusivity of spike-decorate structures using mesoscopic-hydrodynamic simulations. Furthermore, we explored the interplay between affinity and passive viral transport. Our results revealed that the diffusional mechanism of SARS-CoV-2 is strongly influenced by the size and distribution of its spikes. We propose and validate a universal mechanism to explain the link between optimal virion structure and maximal infectivity for many virus families.

## Introduction

Spike-decorated morphologies are ubiquitous in enveloped viruses [1–7]. Depending on the infectious mechanism of the viruses, they exhibit characteristic spikes morphology, number, distribution, and reactivity. In general, the infectivity of a virus is greatly determined by the reactivity between receptor-binding domains (RBDs) present on the spikes and cell receptors (*affinity*), as well as the available number of binding sites (*avidity*). The coronavirus SARS-CoV-2, responsible for the COVID-19 pandemic, is characterized by an ellipsoidal envelope (**E**) decorated with protruding functional *spike* proteins (**S**). In the case of SARS-CoV-2, the alignment and binding of the **S** with the specific angiotensin-converting enzyme-2 (ACE-2) receptor [8] of the human cells determines the linkage and further insertion of the viral genetic material into the cells. Due to its significance, the structural features of the spikes and their effect on the viral infection process have gained increasing attention to streamlining vaccine development and COVID-19 treatment.

SARS-CoV-2 spikes are formed by three protomers of non-covalently bonded protein subunits [9–11]. The binding of the RBD with the epithelium receptor destabilizes the spikes leading to conformational changes in the spike from a tetrahedron-like shape (prefusion) to nail-like (postfusion) morphology [10]. Recent evidence showed that this morphological transition of **S** could also occur before the anchoring to the host cell [8], and suggested that the ratio between pre/postfusion spikes is a hallmark for novel SARS-CoV-2 variants. Investigations on the inactivated strain of the original SARS-CoV-2 from Wuhan (typically denoted as D-form) revealed spikes dominantly in the postfusion state, around 74 percent [10]. In contrast, for mutated variants (also referred to as G-form), only 3% or less were in the postfusion form [11, 12]. Both prefusion and postfusion states appeared randomly distributed on the surface of the envelope. The effect of this morphological transition on the virus infectivity can be significant for vaccine development.

Another distinctive feature of SARS-CoV-2 is a number of spike on the order of *N*_*s*_ ∼ 26 ± 15 per virion [11], where virion corresponds to the complete infective form of the virus outside the host cell. Compared to other viruses, the *N*_*s*_ of SARS-CoV-2 is close to HIV [6] (∼ 14) and murine hepatitis virus [7] (∼ 11), but significantly lower than other enveloped virus such as SARS-CoV [1, 13] (∼ 100), Influenza A [5] (∼ 350), Herpes Simplex [4] (∼ 659), and Lassa [2] (∼ 273), to name a few. SI Table 9. Section 8 has a summary of size and *N*_*s*_ for various common viruses. Considering these differences in *N*_*s*_, an intriguing question is the reason that led the evolution of SARS-CoV-2 towards low *N*_*s*_ and the corresponding effect on virus infectivity. A possible explanation of this disparity is the difference in **S** reactivity towards receptors and antibodies. In principle, from an evolutionary standpoint, a large *N*_*s*_ would favour cell entry and viral propagation. However, this can also lead to an increased vulnerability of the virus as more **S** proteins (epitopes) can be targeted by the immune system [14]. Another justification lay behind the architecture constraints on the maximum *N*_*s*_ a viral envelope can display. Evidence in other coronaviruses [13] has shown that *N*_*s*_ correlates with the size **E** and the flexibility of the constituting membrane proteins. Overall, the reason behind the variability in *N*_*s*_ between enveloped virus and the characteristic lower number for SARS-CoV-2 is yet to be identified. Herein, stemming from a fluid dynamics standpoint, we postulate that spikes morphology, number, and distribution have a crucial effect on virus mobility, which determines the balance between reactivity and transport in ways to promote viral infectivity.

When studying the infectivity mechanism, it is also essential to consider the transport of the virion to the epithelium, where the linkage takes place. The first barrier in the human body against viral infections is the mucus, which covers the surfaces of the respiratory, reproductive, and gastrointestinal tracts [15, 16]. In the mucus, the glycoproteins mucins trap the pathogens forming a gel network with pore size ranging from 200-500nm [17]. However, it has been shown that with such pore size, small virions can diffuse unhindered through the mucus [15] unless the virion form affinity bonds with the mucins. In the respiratory tract, the mucus layer has a thickness from 1 − 10*μ*m [18], and transports the immobilized pathogens to the pharynx for neutralization (see Figure 1.*a*). For SARS-CoV-2, the virion needs to cross that barrier to reach the epithelium before the clearance. As the virions are not self-motile, their motion relies on the advection of the transporting fluid, as well as the diffusion of the virion in the media (see Figure 1.*a*). In general, the diffusive transport set the upper bound on the time scale at which virions can interact with the host cell.

**Figure 1:**
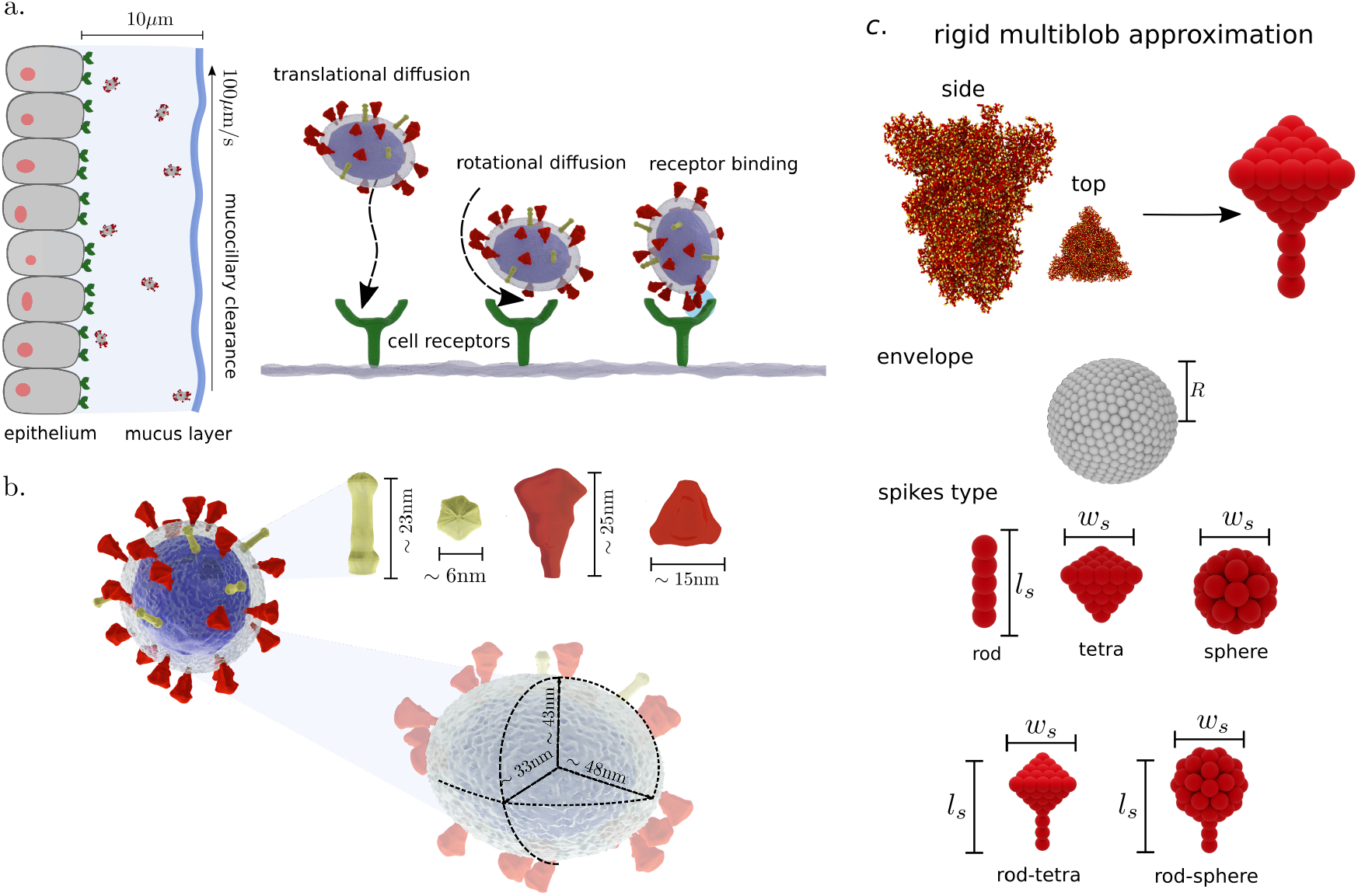
Sketch of viral transport and discretization. **a**. Representation of viral transport within the first barrier of the respiratory system. Virions diffuse while are advected by the mucociliary clearance mechanism (speed ∼ 100 *μ*m*/*s). Effective binding requires translational and rotational diffusion of the virion to the receptor. **b**. Schematic of SARS-CoV-2 characteristic envelope and spikes size. Tetrahedral-shape prefusion **S** in red and needle-shape postfusion **S** in yellow. **c**. Our envelope and spike discretization for viron diffusion based on the size of the spikes reported experimentally. Hydrodynamic interactions on the scale of the spike size are accounted by the model while high frequency atomic motions, fast compared with the viron diffusion, are neglected. Five types of spike morphology are considered: rod, tetra, sphere, rod-tetra, and rod-sphere. Rod, tetra, and sphere are characterized by a single lenght, whereas rod-tetra and rod-sphere require two parameters.

The SARS-CoV-2 virion can be described as a nanosized ellipsoid with radius of their principal axes *R*_1_ ∼ 48nm, *R*_1_ ∼ 43nm, and *R*_1_ ∼ 33nm [11] (see Figure 1.*b*). From the Stokes-Einstein theory, the translational and rotational diffusion arises from the hydrodynamic interactions and depend on the morphological features of the virion and the viscosity of the fluid. Mobility predictions using the Stokes-Einstein theory have shown to be in good agreement with the transport of aquatic viruses [19]. At the nanoscale, the rotational diffusion of such decorated objects may exhibit characteristic deviations compared to a homologous not-functionalized ellipsoid. Furthermore, since the shape and distribution of **S** can alter its diffusion rate, a detailed characterization of the virions diffusivity may provide rheological signatures to differentiate various virus types, as shown in the pioneer work by Kanso and coauthors [20, 21] using rigid bead-rod theory.

Here, we adopt the rigid multiblob methodology [22] (RMB) (see Method section and Section 1. SI) to investigate the mobility of virion models computationally. Using the recently reported structure of SARS-CoV-2 [11], we consider the effects of spikes distribution and morphology on the complex diffusive transport of the virus suspended in a single fluid. Using the RMB, we construct precise mesoscale models that allow us to incorporate consistently thermal fluctuation effects and hydrodynamic interactions with the suspending media. In these models, we neglect **S** mobility on the envelope surface. However, the approximation allows us to elucidate the morphological features that affect the hydrodynamic behaviour of the SARS-CoV-2 and offer potential applications as rheological biomarkers. Moreover, we show that the hydrodynamic characterization elucidates a universal mobility mechanism among different families of enveloped viruses.

### Virion Diffusion

Using the RMB we discretize the virion as a set of rigidly connected blobs (see Figure 1.*c*), and consider that it moves as a rigid object. We construct virion models with spherical and ellipsoidal **E** of size *R*, and investigate the effects of spikes shape on mobility using five different spikes inspired by the morphologies reported in the literature for various viruses (i.e. HIV, MVH, Denge, SARS-CoV, Lassa, Herpes, Influenza). We adopt the following labelling for the investigated shapes: *rod, sphere, tetra, rod-sphere*, and *rod-tetra*. Rod, tetra, and sphere shapes posess only one characteristic size: length (*l*_*s*_*/R*) or width (*w*_*s*_*/R*), whereas for rod-tetra and rod-sphere both *l*_*s*_*/R* and *w*_*s*_*/R* are defined (see SI Section 7, Figures 4 and 5). To account for **S** distribution on the surface of **E**, we construct virions with both homogeneous and randomly localized **S**. For SARS-CoV-2 [11] a realistic representation is constructed using an ellipsoidal **E**, with rod-tetra (prefusion) and rod (postfusion) **S**, localized in random configurations.

Based on the currently known structures [11] of SARS-CoV-2 in their original (D-form) and mutated (G-form) strains, we give, in Table 1, a breakdown of the numerically-estimated diffusion coefficients for three different media viscosities: water, blood, and mucus (see SI.Table 12 a full list of *D*_*t*_ and *D*_*r*_ for other virions). Strictly speaking, the diffusion of the virion in complex fluids is strongly determined by microrheological features such as mesh size (mucus) or the presence of other constituents on the length scales of the virion. Given the size of SARS-CoV-2 (∼ 90nm), we expect it to diffuse nearly unhindered on typical mucus meshes (200-500nm). However, we also provide the estimated diffusion coefficients using the macroscopic viscosities of blood and mucus as an upper-bound indicative for the virion diffusivity. Diffusion coefficients in crowded biological environments have been reported to decrease over 1000-fold, compared to diffusion in water [23]. For comparison, in SI Table 4 and 5, we compiled the estimated diffusivities of SARS-CoV-2 on the range of the variance on number of spikes reported.

**Table 1:** Translational and rotational diffusivities computed for different virions in water, blood, and nasal mucus. The diffusivities of the virions are computed using equation (5) in Methods section. The viscosity *η* values are for water = 0.001 Pa · s, blood = 0.0035 Pa · s. [24] and, nasal mucus = 1.6 Pa · s [16]. Temperature = 298.15 K, and *k*_*b*_ = 1.38064852 · 10^−23^ m^2^kg*/*s^2^K.

| Virus | water |  | blood |  | nasal mucus |  |
| --- | --- | --- | --- | --- | --- | --- |
| | $D_t [\mu\text{m}^2/\text{s}]$ | $D_r [1/\text{s}]$ | $D_t [\mu\text{m}^2/\text{s}]$ | $D_r [1/\text{s}]$ | $D_t [\mu\text{m}^2/\text{s}]$ | $D_r [1/\text{s}]$ |
| SARS-CoV-2 (G-form) | 4.0451 | 965.4694 | 1.1557 | 275.8484 | $2.5281 \cdot 10^{-3}$ | 0.6034 |
| SARS-CoV-2 (D-form) | 4.3345 | 1185.2682 | 1.2414 | 338.6481 | $2.7156 \cdot 10^{-3}$ | 0.7408 |

In the remaining, to streamline the discussion, we introduce a reduced translational 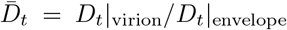 and rotational 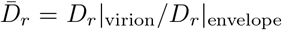 diffusion coefficients. Where *D*|_envelope_ is the calculated diffusivity for an envelope without spikes. Reduced diffusivities allow us to rationalize the results in terms of spikes count and morphology. The results described herein correspond to resolutions with discretization errors below 3%. while convergence results are presented in the Section 4 of the supporting information (SI Table 1-5 summarize the convergence errors for the **E** optimal **E** resolution, resolution convergence for the whole SARS-CoV-2 virion **E** and **S** and covergence errors for tetrahedral **S** respectively)

### Envelope shape and spike distribution

In general, axial asymmetry of **E** can favour directional motion of the virions [18]. However, for SARS-CoV-2 **E** we observed weak effect of envelope shape on virion mobility, only on the order of 1% for 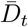 and 2% for 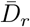 (see 2.*a*). The later, may indicate that the dominant effect of ellipsoidal shapes is to maximize the available surface area for the spikes in **E**, whereas keeping virion transport unaffected. In contrast, the distribution of **S** showed to have a more important role on virion transport. For all the spike shapes investigated we found a relative increment on both 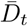 and 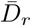, when spikes are localized at random positions around the surface relative to the homogeneous case (see SI Section 6 S.Figure 3 and S.Table 8). In particular, for rod-tetra type (used for SARS-CoV-2), the increment on 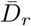 ranged from 10% to 3%. We speculate that the sparsity and randomness of **S** favour the ability of the virion to explore its surroundings to reach binding receptors. This is consistent with all-atoms simulations [25] of **S**, which suggested that the conformational freedom of the spikes in **E** may increase the infectivity of the virus by providing mechanical robustness, facilitating motions to avoid antibodies access, and increasing the avidity when binding the cell [25].

### Spikes shape and size

In Figure 2.*a*, we compare the reduced translation and rotational diffusivity for the five types of spikes studied for virions with 12 homogeneously distributed spikes, and fixed spike size, for both spherical and ellipsoidal envelopes. The presence of spikes induces a reduction in the translational diffusion of the virions between 20 to 30 per cent (compared to the naked envelope), whereas the impact in rotational diffusion is more significant, ranging from 50 to 70 per cent. Significantly, the spike shape determines the extent of the reduction on both 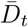 and 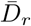. Overall, larger spikes affect the transport properties of the virion strongly. The small but noticeable differences in the diffusion between globular and tetrahedral spikes indicate a characteristic transport signature that can be further exploited for virus identification. In SI Section 13 (S.Figure 15 and 16), we show the relative differences in mobility for the **S** shapes evaluated.

**Figure 2:**
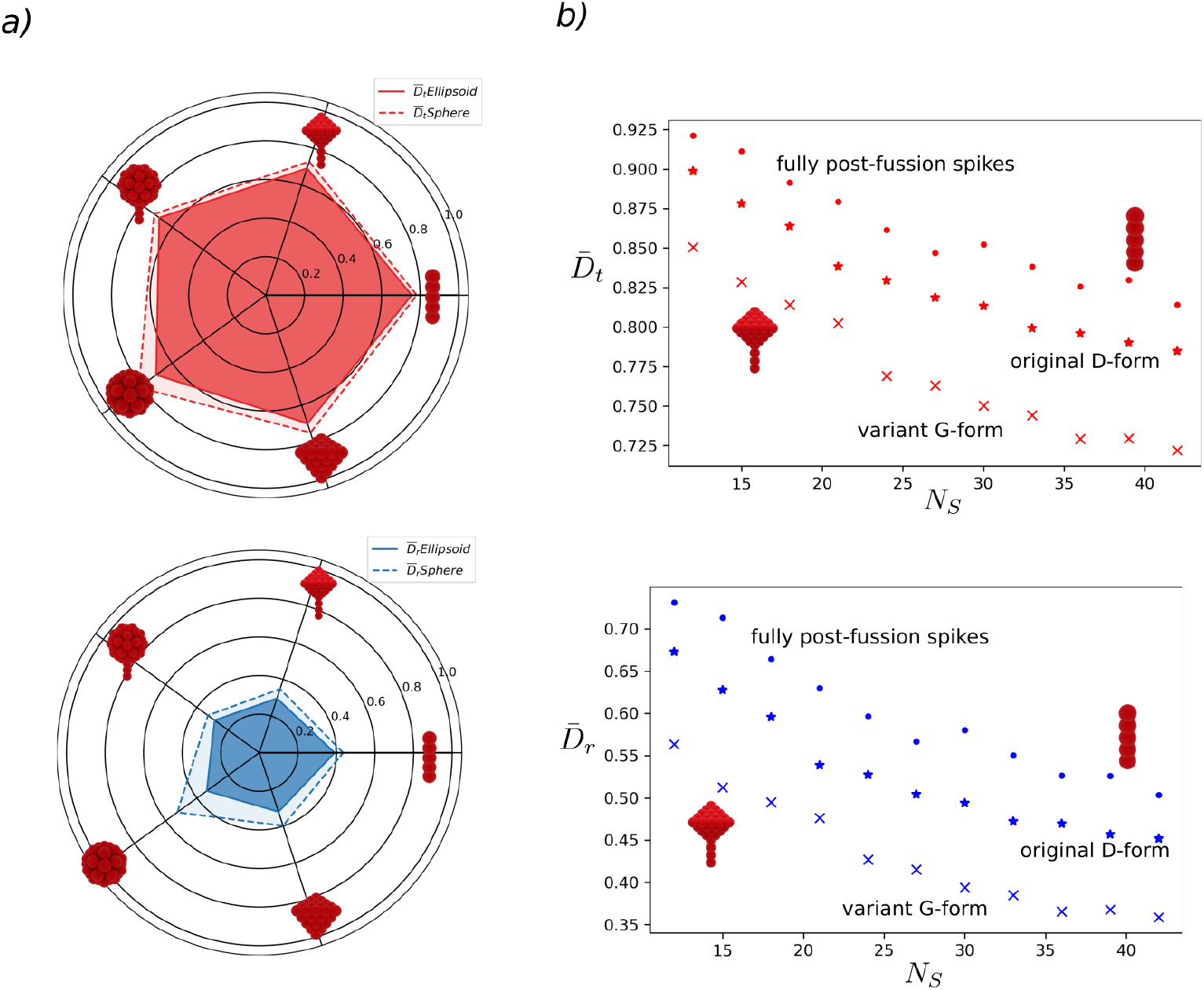
Effect of **S** morphology on the mobility of virions. **a**. Comparison of the reduced translational and rotational diffusion for the different **S** shapes for virions with *N*_*s*_ = 12 spikes, and *l*_*s*_*/R* = 0.4, homogeneously distributed. Ellipsoidal envelopes exhibit slightly larger deviations on the rotational diffusion than spherical envelopes due to their small asymmetry. The differences in 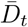 between **S** shapes do not reveal a significant difference. In contrast, for 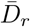, small but observable differences indicate the potential use of the rotational diffusion of virions as a rheological biomarker. **b**. Variation on 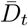 and 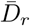 with *N*_*s*_ for fully postfusion (rod), fully prefusion (rod-tetra) and mixed postfusion/prefusion **S**. The diffusion coefficient values for each *N*_*s*_ are obtained from ten independent realizations with **S** randomly distributed on the envelope. The fraction of postfusion **S** in the mixed case corresponds to the original D-forms of SARS-CoV-2, *N*_*s*_|_post_*/N*_*s*_ = 0.7. Fully prefusion case is consistent with mutated G-forms characterized by *N*_*s*_|_pre_*/N*_*s*_ ∼ 1. Regardless of the differences in **S** morphology, the reduction in both 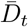 and 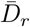 exhibits the same functional dependence as *N*_*s*_ increases. However, the magnitude of the mobility reduction is larger for the full-prefusion case.

Regarding the effect of the size of the spike (*l*_*s*_*/R* and/or *w*_*s*_*/R*) we found, as expected, that larger spikes considerably reduce the diffusion of the virions in all cases. Nevertheless, the dimensionality of the spikes affects the scaling of diffusion with the spike size. For instance, comparing rod- and tetra-type shapes, rod shapes (that are dominantly one dimensional) showed a weaker variation on 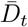 and 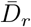 as the size increases (see SI Figure 4). In contrast, tetra-shape **S** displayed a strong reduction on diffusivity (see SI Figure 5). Overall, we observed that the virions with bulkier and larger spikes (compared to the envelope size) had an intrinsic diffusional penalty. Therefore, the regulation on the number of spikes suggests a possible alternative to compensate for this reduction in mobility.

### Number of spikes

Now, we switch attention to the role of the number of spikes on the mobility of the virion. In Figure 2.*b*, we compile the variation on translational and rotational diffusion for virions with different rod and rod-tetra shape spikes. Overall, leading to reductions ranging from 10 to 30 percent in the 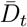 and 30 to 70 percent in 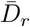, as the number of spikes increases. Interestingly, the dependency of 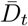 and 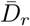 with the number of spikes decays in the same fashion, regardless of the spike type. In Figure 2.*b* the range *N*_*s*_ presented corresponds to the experimentally reported for SARS-CoV-2 virions with fully-postfusion **S** (rod), fully-prefusion **S** (rod-tetra), and mixed **S**. The later is consistent with the original Wuhan strain, or D-form. Whereas fully-prefusion resembles mutated strains, G-form. In general, the larger population of **S** in prefusion states in mutated variants induces a reduction in the mobility of the virion, and this decay occurs quickly after a few number of spikes are added on the surface of **E**. Interestingly, the mobility appears to reach a saturation condition where increasing *N*_*s*_ no longer significantly affect the diffusion rate of the virions.

To explore further the decay in mobility due to *N*_*s*_, we modelled various enveloped viruses (Figure 3.*a*) using the characteristic sizes reported in the literature (in SI Table 10 we compile *N*_*s*_, size, and **S** shape for de different virions, and in Tables 11-12 the estimated reduced difusivities). Remarkably, we identified that the form of the decay in 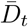 and 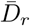 (see SI Figure 6 and 7) is consistent in other viruses, as shown for 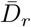 in Figure 3.*a*. However, the magnitude of the drop in mobility depends on the **S** shape and size. For example, Herpes virions with *l*_*s*_*/R* = 0.22 can exhibit upto 23% drop in 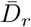, whereas SARS-CoV-2 with *l*_*s*_*/R* = 0.52 showed reduction up to 70%.

**Figure 3:**
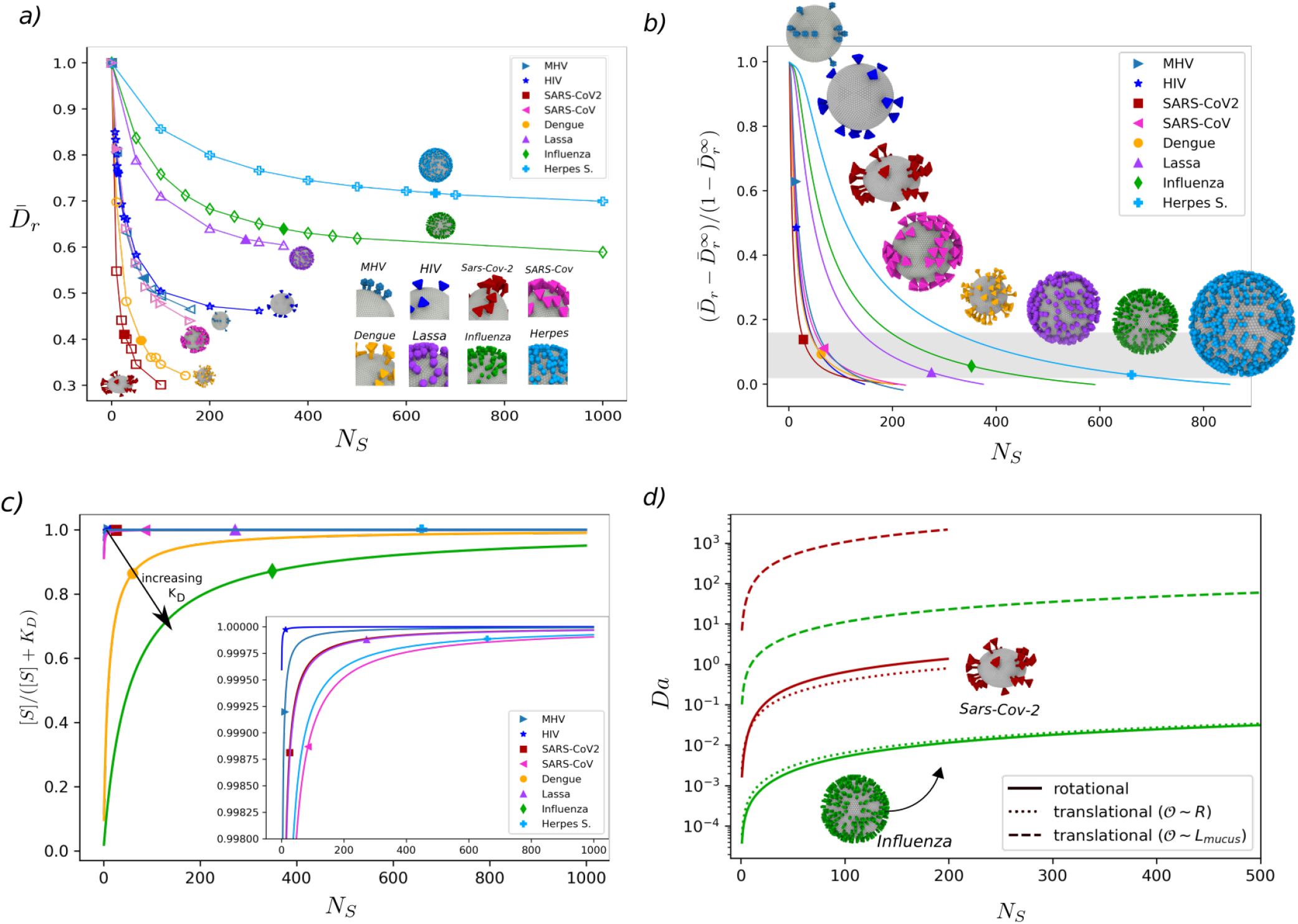
Interplay between virion rotational diffusion and receptor binding affinity for SARS-CoV-2 and selected viruses. **a**. Effect of *N*_*s*_ on the rotational diffusion. Each virion family exhibits a characteristic reduction on its mobility reaching an asymptotic value for large *N*_*s*_. SARS-CoV-2 shows the sharpest drop in diffusion, whereas for Herpes Simplex the reduction occurs over a much larger *N*_*s*_ range. Inset: Snapshots of the morphologies used to represent each virion type. In the Methods Section we include the reported crystal structure and sizes of the spikes for comparison. **b**. Excess of rotational diffusion vs. *N*_*s*_ fitted from Eq. (1). The fitting parameters are summarized in Table 2 and SI Section 9. Filled markers indicate the experimentally reported *N*_*s*_ for the different viruses. Viruses share an equivalent diffusional state independently of *N*_*s*_. This ground state is likely induced as a balance between their geometrical features and spikes reactivity. The higher Δ*D*_*r*_ in HIV and MHV can be explained due to an enhanced mobility of **S** on the **E** surface. Such effect is not currently accounted in our model. **c**. Saturation function change with *N*_*s*_ for various virus based on their binding constant *K*_*D*_ to cellular receptors. The magnitudes of *K*_*D*_ are summarized in SI Table 14. The majority of the viruses posses a high saturation value (∼ 1) over the range of *N*_*s*_ reported. Influenza A and Dengue show the lowest level of saturation. This lower saturation for Influenza A viruses may explain its characteristic variety of pleiomorphic structures [5] as a way to enhance infectivity. Inset plot: Zoom over the saturation levels of SARS-CoV-2, HIV, Herpes and Lassa virions. HIV reaches saturation at very low spike numbers. **d**. Translational and Rotational Damkohler number change with *N*_*s*_for SARS-CoV-2 and Influenza A virus. At the lenght scales of the **E** size *R*, the reaction time for Influenza viruses is kinetically controlled due to its low affinity, whereas for SARS-CoV-2 the rotational and kinetic time scales are on the same order. When accounting for the whole binding process, the translational diffusion on the lenght scales a mucus layer with thickness of *L*_mucus_ = 5 *μ*m amounts for the larger part of the required time.

Considering that the diffusivity of the virions varies between one corresponding to an envelope without spikes 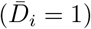 and one with closely packed spikes 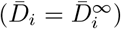 we introduce the *excess* diffusion coefficients 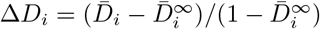, for *i* = *t, r*. This expression varies from 1 when *N*_*s*_ = 0 to 0 when 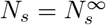. The term 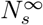 is the number of **S** at which the virion mobility saturates, 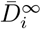, and the effect of *N*_*s*_ is negligible. Based on the computed translational and rotational diffusivities we postulate the following expression to describe the excess diffusion dependency with *N*_*s*_

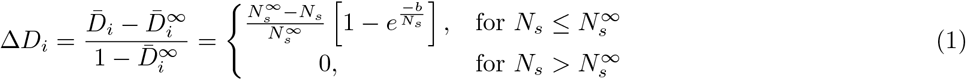

where 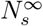 and *b* are fitting parameters that depend on the characteristic size of the spike. The first term on the right-hand side of (1) accounts for a linear dependency on the number of spikes before reaching 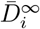. Similar linear dependence on *N*_*s*_ has been identified for ligand-receptor interactions using functionalized colloids [26]. The second term on the right-hand side of (1) describes an exponential decay in the diffusivity that is controlled by the shape-dependent parameter, *b*. Using the calculated 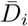 and taking 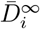 from the largest number of spikes simulated, equation (1) allows us to obtain the characteristic values of 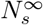 and *b*, for various families of enveloped virions as presented in Table 2 and SI (Figure 8 and Table 13 Section 9). In Figure 3.*b*, we depict the variation in Δ*D*_*r*_ for different viruses along with the experimentally reported *N*_*s*_. Except for HIV and MHV (with Δ*D*_*r*_ ∼ 0.5), independently of the type, we identify a characteristic trend in the mobility, with a mean Δ*D*_*t*_ ∼ 0.12 and Δ*D*_*r*_ ∼ 0.08 among all the viruses. This suggests the existence of a general transport mechanism across enveloped viruses. We speculate that this mobility regime coincides with condition where reactivity and mobility balance out. For each type of virion, the shape and size of **S** determines hydrodynamically how much the mobility can change, while the reactivity of **S** sets the extent of such reduction in 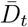 and 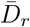. Thus, this interplay is conserved across virus families. In the following section we address this hypothesis.

**Table 2:**
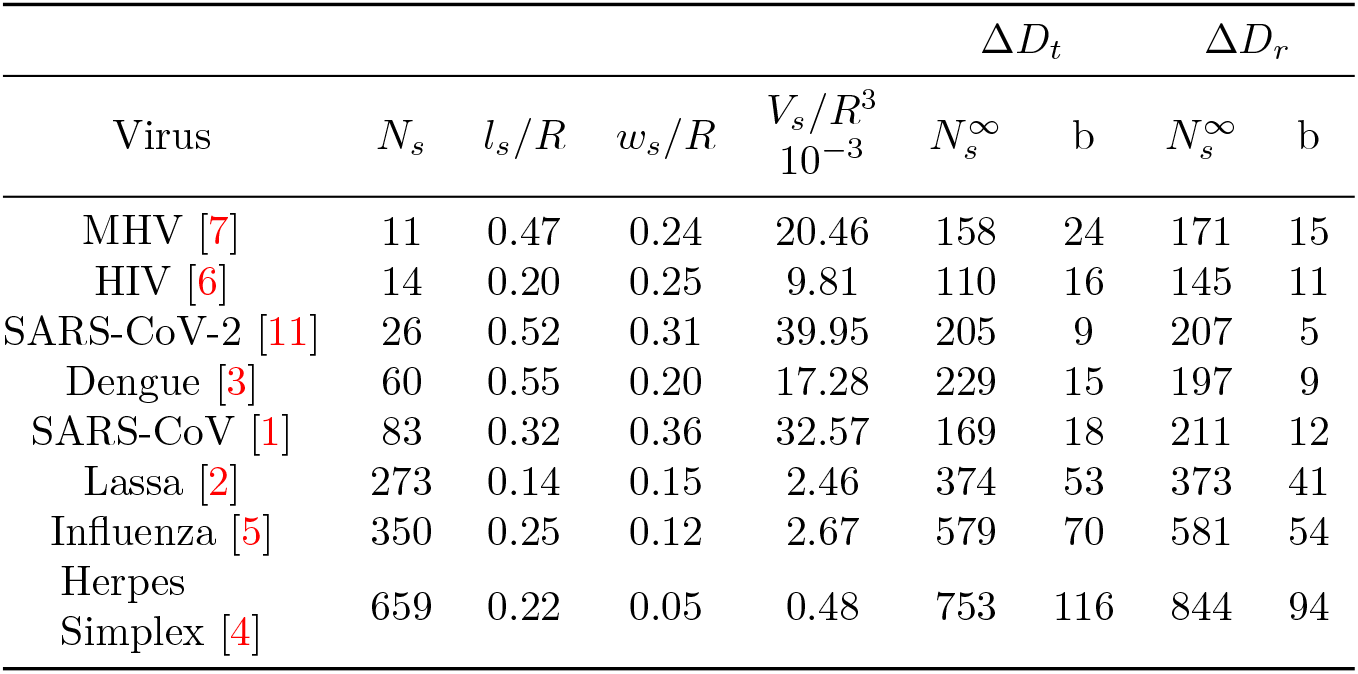
Summary of characteristic *N*_*s*_, *l*_*s*_, *w*_*s*_, and spike volume *V*_*s*_*/R*^3^ for different virions, along with the fitted parameters 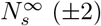 and *b* (±2 for Δ*D*_*t*_ and ±3 for Δ*D*_*r*_) from (1). All the parameters obtained from fitting of (1) lead to a determination coefficient 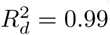.

### Affinity-Mobility balance

As the viral spikes of SARS-CoV-2 mutate into new variants with potentially different affinities but equivalent spikes size, it is necessary to elucidate the relative balance between affinity and mobility. To consider only affinity effects, it is customary to determine the strength of the spike-receptor interactions measuring the binding affinity constant, *K*_*D*_ [27]. For a given *K*_*D*_ and concentration of spikes [*S*] surrounding the envelope, it is possible to introduce a saturation function [*S*]*/*([*S*] + *K*_*D*_) that relates the affinity with the number of spikes. The spike concentration [*S*] variation with *N*_*s*_ is approximated using the volume around the envelope. This volume is given by the radius of the envelope and the height of the **S** (see SI Section 10). The saturation function varies from 0 ([*S*] *<< K*_*D*_) when the concentration of spikes is a limiting factor for binding to occur, to 1 ([*S*] >> *K*_*D*_) when spikes availability is ensured facilitating binding. In Figure 3.*c*, we present the variation of the saturation function for G-forms of SARS-CoV-2, along with another viruses for comparison (see SI Table 14 for the *K*_*D*_ values of the viruses and receptors). For SARS-CoV-2 HIV, Lassa, and Herpes, we find that the saturation function reaches values close to unity for a small number of spikes. In contrast, Influenza A exhibits the lowest saturation, even at significant *N*_*s*_. In general, after comparing the experimentally measured *N*_*s*_, the effect of *K*_*D*_ alone is not sufficient to rationalize the difference in *N*_*s*_ among the investigated virions.

Since the effective reaction time *t*_*r*_ for virion/receptor association would depend on diffusional *t*_*d*_ and binding *t*_*b*_ time scales, we use the non-dimensional Damkohler number Da = *t*_*d*_*/t*_*b*_, to identify the controlling mechanism virion/receptor association. The diffusional time *t*_*d*_ = *t*_*t*_ + *t*_*r*_ accounts for both translation and rotation leading to the definition of both Da_*t*_ and Da_*r*_. In general, *t*_*b*_ controls the reaction rate for low-affinity interactions, whereas *t*_*d*_ determine reaction rate for interactions requiring the alignment or localization of the ligan/receptor pair [28]. Theoretical models on ligand-receptor interactions mediated by rotational diffusion [26] showed that *t*_*r*_ is linear function of the *N*_*s*_ and the relative surface occupied by **S** [26]. Here, we estimate *t*_*b*_ using the reported asociation reaction constant *K*_*on*_ between **S** and ACE-2 [8, 29], for the original D-form (*K*_*on*_ = 1.37 · 10^5^ [Ms]^−1^) and mutated G-form (*K*_*on*_ = 7.92 · 10^4^ [Ms]^−1^) SARS-CoV-2 strains. For SARS-CoV-2, we obtain Da_*r*_ ∼ 0.4 and Da_*r*_ ∼ 0.2 for the D- and G-forms, respectively. Similar order of magnitude is observed for Da_*t*_ for diffusion on the lengh scales of the virion size. These values of Da ∼ 1 evidence that SARS-CoV-2 and its variants are effectively on a regime where both affinities and mobilities are highly coupled. This value is in contrast with the affinity-controlled regime of Influenza A that exhibit the lowest affinity among the virus investigated (see SI Table 14) with *K*_*on*_ = 700 [Ms]^−1^ [18], leading to Da_*r*_ ∼ 1 · 10^−2^. In Figure 3.*d*, we present the variation of Da_*t*_ and Da_*r*_ for different spikes number for SARS-CoV-2 and Influenza A, for comparison. For the first entry, considering that the virion crosses a mucus barrier of ≈ 5*μ*m the maximum diffusional time 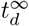 is determined by 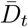 over the length scale of the mucus thickness. Thus leading an overall binding process dominated by transport Da > 1. In SI Section 12 (Figures 12-14), we have summarized translational and rotational time scales for the different virion types, for the first entry stage.

If we consider that the energy of binding between RBDs and a cell receptor is *ϵ*_bind_, the strength of all the interactions combined (upto a first order approximation) can be expressed as Γ = *N*_*s*_*ϵ*_bind_. In general, increasing the number of spikes should favor virion avidity, whereas reducing the diffusion rate of the virion as shown in Figure 3.*b*. For convenience, we introduce a geometrically constraint avidity 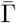, given by

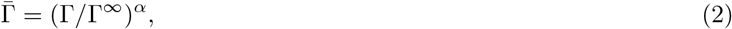

such that 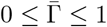, independently of the value of *ϵ*_bind_. The exponent *α* in (2) incorporates no-linearities in the binding between RBDs and a cell receptors. For linearly dependent interactions we have that *α* = 1. The limit 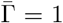 indicates the saturation condition, where the increase in *N*_*s*_ no longer influences the binding. The magnitude of 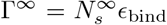 depends on the characteristic envelope size *R* and spike volume. The use of 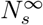 to determine the behavior of 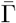 assumes that both mobility and reactivity saturate on the same order of *N*_*s*_, however, this may change as the order of reaction between **S** and receptors changes.

Now, we define an infectivity parameter 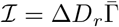 that considers the interplay between the transport and reactivity. On one hand, independently of the strength of the interactions between the virions and the epithelium, the time required for the virion to reach available anchoring sites is inversely proportional to the diffusivity of the virus. On the other hand, regardless of the time taken to the virion to reach the receptors, a succesful binding depends on the avidity of sites. The parameter ℐ accounts for the transport limiting condition through Δ*D*_*r*_, and the reaction limiting condition through 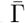. In Figure 4, we present the variation of ℐ with *N*_*s*_ for SARS-CoV-2. Remarkably, the maximum in ℐ that balance avidity and mobility coincides with the reported values for spikes count for SARS-CoV-2 [11] (26±15). Similarly, the maximum in the infectivity curve for other virion models SARS-Cov, Lassa, Denge, Herpes Simplex, and Influenza A is in close agreement with experimental evidence [1–5]. In Figure 4 we use a corrected exponent *α* that depends on the available area of **S**. The corrected exponent is given by *α* = 1 + *b/*100, where *b* is the shape-dependent parameter obtained from (1). We remark that HIV [6] and MHV [7] showed a small deviation from the maximum ℐ. The differenciating behavior for HIV, can be related with the presence of highly reactive **S** able to reach early avidity saturation, Γ^∞^, or the ability of **S** to diffuse on the **E** surface [30]. These effects are not accounted in our current model, however, they will be addressed in future publications.

**Figure 4:**
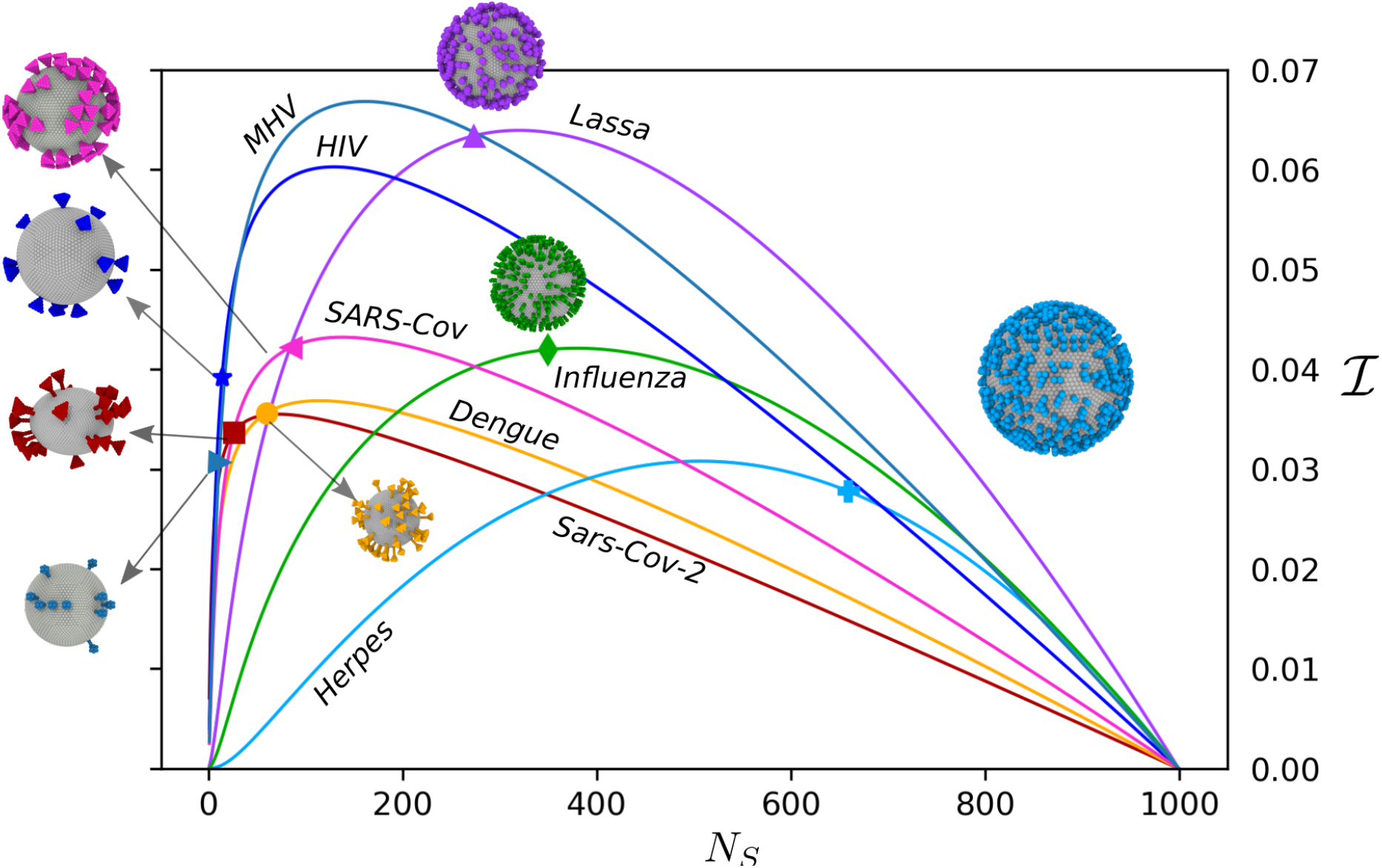
Interplay between virion rotational diffusion and receptor binding affinity for SARS-CoV-2 and selected viruses. The figure shows the variation in the infectivity parameter ℐ with the number of spikes. The markers indicate the experimentally reported values of *N*_*s*_ for each virion type, which in most cases agree well with the theoretical optimum value. HIV and MHV are two exception with particular low values of *N*_*s*_, perhaps related with enhanced mobility of the spikes around the envelope, or an earlier reactive saturation requiring fewer **S** than 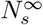 due to strong binding affinities with their receptors.

## Discussion

The emergence of novel SARS-CoV-2 variants poses a significant challenge. Important information is known regarding the location and effect of the existent mutations [27, 29], providing relevant insights. However, the transmissibility of viruses within and among hosts may exhibit features that go beyond the particular genetic sequence [18]. In this context, morphological features can be studied to reveal similarities and differences between different virus families. The current results provide an initial step for further microrheological characterization of viral solutions that can serve as a tool to identify potential biomarkers and overall gaining an understanding of viral functionality.

We show how the interplay between spikes distribution, shape, size affect the mobility of the virions. We postulate that transport properties of the virions roots from geometrical constraints that can explain the differences in spikes density across a variety of virus. Thus, these geometrical constraints, along with the affinity of the RBD of the spikes, may indicate how different virus families exist on optimal evolutionary conditions. The saturation values on **S** affinity and along with ℐ, justify the lower *N*_*s*_ reported for SARS-CoV-2 and its variants. From an evolutionary standpoint, SARS-CoV-2 may have reached an optimal avidity/mobility balance that ensures a large mobility due to a moderate value of *N*_*s*_ and a large affinity to the receptors groups thanks to a high saturation value.

The understanding of the virus spreading through the first barriers of defense in our body is in general difficult, the complexity and nonlinearity of the interactions between the media and viruses makes difficult the investigations using experiments alone. Our results offer a good approximation of virion transport properties based on physiological conditions that can be further used in viral modelling. Additionally, in a more general sense can potentially guide the design of vectors for nasal vaccines that optimize immune response [31]. Our findings also provide tools for the designing of microrheological devices for screening, detection, and characterization of viruses.

## Supporting information

Supporting Information: Method description and calibration. Reported data for various virus.

## Acknowledgments

This research is supported by the Basque Government through the BERC 2018-2021 programme and by the Spanish State Research Agency through BCAM Severo Ochoa excellence accreditation SEV-2017-0718 and through the project PID2020-117080RB-C55 funded by (AEI/FEDER, UE) with acronym COMPU-NANO-HYDRO. The authors acknowledge also the financial support received by the Basque Business Develop ment Agency under ELKARTEK 2019 programme (bmG19 project: grant KK-2019/00015) and through the “Mathematical Modeling Applied to Health” Project. N.M acknowledges the support from the European Union’s Horizon 2020 under the Marie Skłodowska-Curie Individual Fellowships grant 101021893, with acronym ViBRheo. F.B.U. acknowledges support from “la Caixa” Foundation (ID 100010434), fellowship LCF/BQ/PI20/11760014, and from the European Union’s Horizon 2020 research and innovation programme under the Marie Sklodowska-Curie grant agreement No 847648.

## Methods

### Virion diffusivity

At low Reynolds number the deterministic motion of the virion can be approximated in terms of the Stokes equations, such that

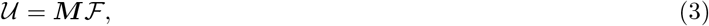

where 𝒰 ={**u**, *ω*} is the vector of the linear (**u**) and angular (*ω*) velocities, ℱ = {**f**, *τ*} is the vector formed by the total forces (**f**) and torques (*τ*) exerted on the body. The tensor ***M*** has denoted the mobility and depends only on the shape of the body. This tensor provides information about the hydrodynamic interactions acting on the body. Here, we determine mobility ***M*** by solving the Stokes equations with the rigid multiblob method [22] (see SI Section 1. for a detailed description). The virions envelope and spikes are discretized as a set of rigidly-connected blobs of size *r*_*b*_, located at a distance *r*_*o*_ = 2*r*_*b*_ between blob centers. Depending on the size of the modelled object, *R*_object_, and the distance between blobs, *r*_*o*_, we define the resolution as *R*_object_*/r*_*o*_. In SI Figure 1 and 2 we illustrate the characteristic discretization dimensions of **E** and **S** respectively. In Section 11 of SI we describe the construction of discrete morphologies for **E** (SI Figure 9) and **S** (SI Figure 10 and 11). We use resolutions fine enough to compute the mobilities with errors below 3%, see SI Section 4 for convergence results.

The translational [32] and rotational [33] diffusivity of can be then computed using the numerical approximation of ***M***, as

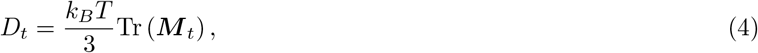

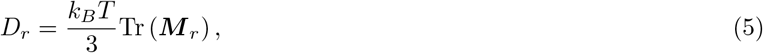

Where *k*_*B*_ is the Boltzmann constant, *T* is the temperature. The tensors ***M*** _*t*_ and ***M*** _*r*_ are the translational and rotational components of the mobility. This approximation considers that the virus moves as a rigid object, thus neglecting spikes mobility on the envelope surface. Nevertheless, this approximation allows us to elucidate the morphological features that affect the diffusion of the virions. Such morphological features offer potential applications further to analyze similarities and differences between different virus families.

### Envelope shape

Virion shape and size have been reported to affect the efficiency of the virus to penetrate through mucus mesh (the first barrier before accessing endothelial cells in the upper airways). For influenza, for instance, pleiomorphic variations have been speculated to correlate with infectivity and pathogenicity [5]. In the classic cartoon of SARS-CoV-2, it is considered as a spherical envelope surrounded by tightly packed and homogeneously distributed spikes. However, recent investigations using cryoelectron tomography (cryo-ET) and subtomogram averaging (STA) provide us with a realistic picture of the complete three-dimensional description of SARS-CoV-2 and its main morphological features [11]. Given the ellipsoidal morphology of SARS-CoV-2, we compare both spherical and ellipsoid-shape envelopes to provide a better description of SARS-CoV-2 motion while providing the foundation for transport of another spherical-shape virus.

Spherical and ellipsoidal solids exhibit differences in their translational and rotational diffusion as the aspect ratio between the principal axis changes. For spheres, the Stokes-Einstein equation provides a direct correlation of both *D*_*t*_ and *D*_*r*_, which is consistent with the spherical envelop model that we use herein. For ellipsoids, we validate the envelope model in terms of its average diffusivity *D*_ellip_ = 1*/*3 Σ*D*_*i*_ (for *i* = *x, y, z*), with respect to the diffusivity 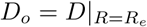, of an equivalent sphere with the same volume and equivalent radius *R*_*e*_ (see SI Section 4.1 for detailed description of *R*_*e*_). For the SARS-CoV-2 ellipsoidal envelopes investigated, with ratios between the minor and major radius of the principal axes of 0.9 and 0.7, we obtain *D*_ellip_*/D*_*o*_ ∼ 0.98. This is consistent with semianalytical derivations of drag coefficients [34] that lead to ratios on the order of 0.99.

### Spikes shape

To investigate the effect of shape, we construct a model of spikes consistent with the morphologies presented in Figure 1.c. In Figure 5, we present the reported morphology and size for the different virions investigated. SI Table 10 compiles the characteristic sizes of the corresponding discretized virions in RMB. These shapes correspond to coarse representations of **S**, able to capture hydrodynamic interactions on the scale of the spike size. Finer representations, including high-frequency sub-nanoscopic details, are hindered by the overall motion of the virus, and their effect on the measured diffusivity are below the resolution of the model. Herein, the finest resolution is selected to ensure accurate hydrodynamic modelling of the different spikes morphologies.

**Figure 5:**
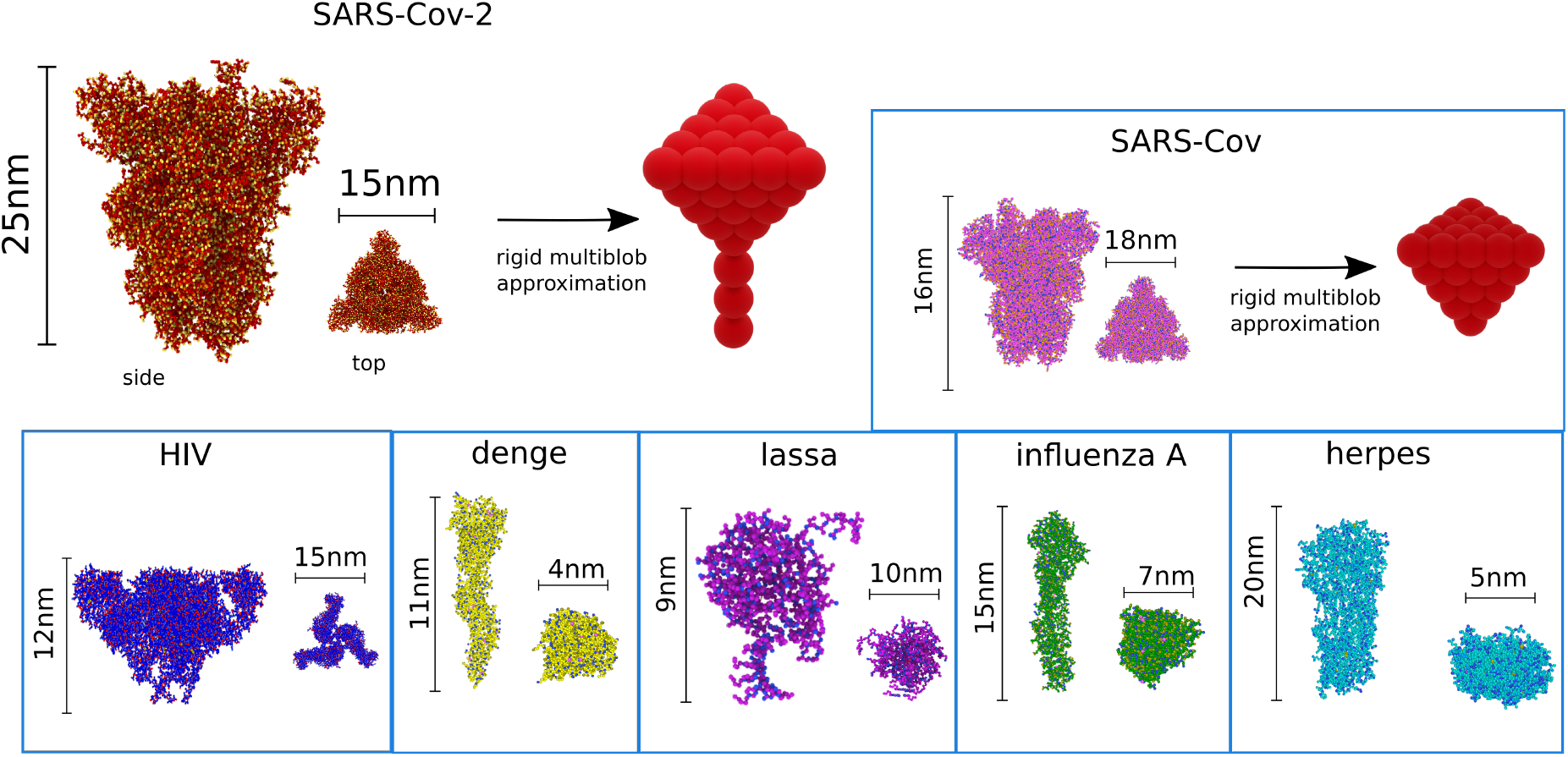
Discretization adopted to model the spike morphology of different virions. The characteristic sizes are shown for: HIV [35], dengue [36], SARS-Cov-2 [37], SARS-Cov [38],lassa [39], influenza [40], and herpes [41]. The basic set of discrete morphologies are rod, tetra, sphere, rod-tetra, and rod-sphere.

Previous investigations have shown differences in the estimated rotational diffusion between single [20] and three [21] beaded representations SARS-CoV-2 spikes. Even though three-beaded models approximate better the tetrahedral shape of the SARS-CoV-2 spikes, the minimal accurate spike representations are still to be determined. To rule out effects due to discretization, we test various ratios between spike size *l*_s_, and inter-blob distance *r*_*o*_. In SI Section 4.3 (Table 3) and Section 4.4, we present the variation on the measured diffusivity for different spike resolutions. For practical purposes, we identify that *l*_s_*/r*_*o*_ > 5 is enough to describe within 1% error both translation and rotation of spherical objects, whereas, for tetrahedral shapes, the associated error is 8%. However, further refinement of the tetrahedral spikes allows us to reduce the error to ∼ 1% (see SI Section 4.4, Table 4-5 and Figure 2). In Table 3 we present the computed 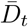 and 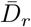 for SARS-CoV-2 using different resolutions. We define the error in our approximation considering the finest (most accurate and computationally expensive) resolution modelled. Coarser representations (four blobs per **S**) with resolutions of *R/r*_*o*_ = 3.6 induced errors on the order of 20% for 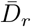. Here, we adopt a resolution of 14.5 leading to a maximum numerical error of 0.7%.

**Table 3:**
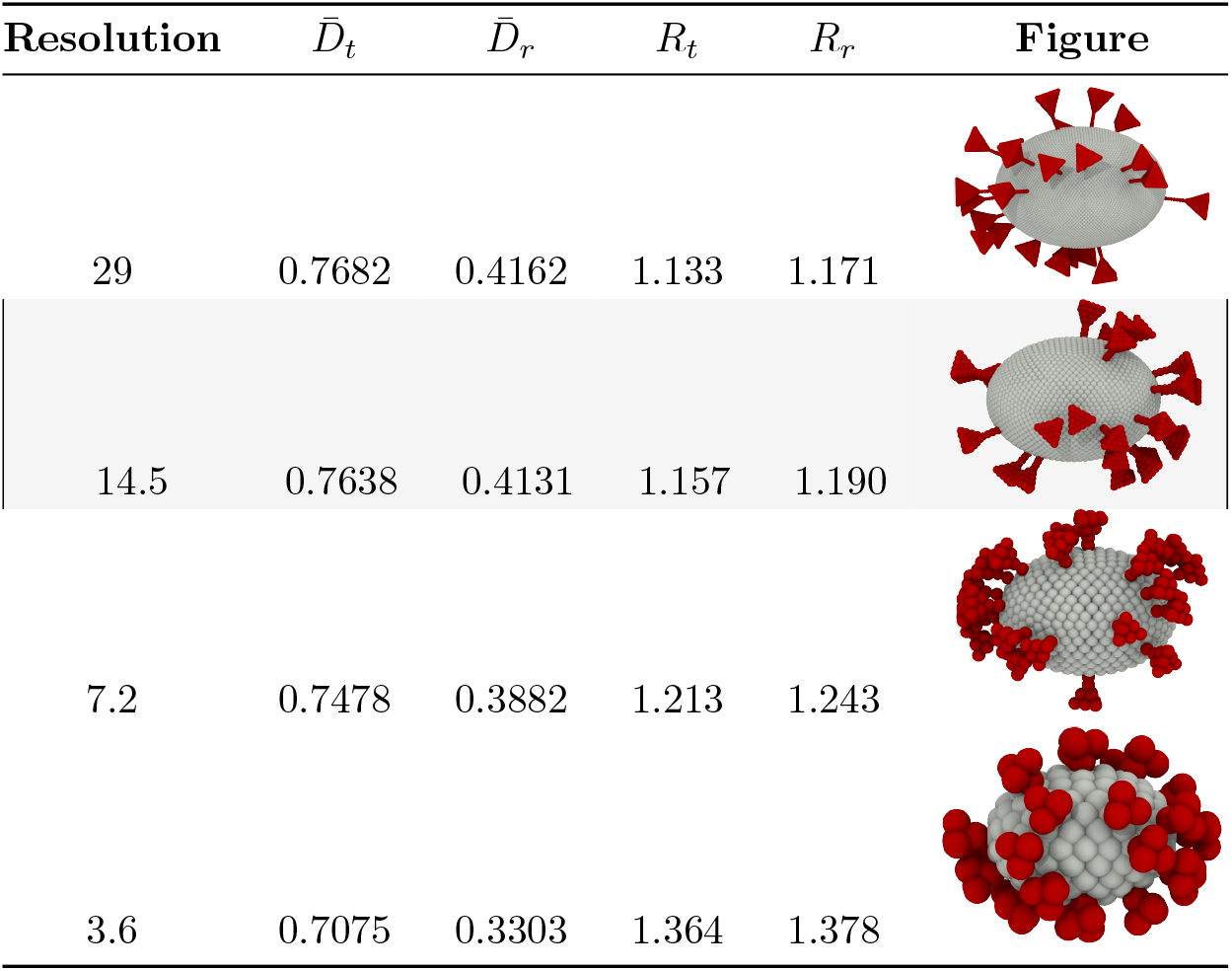
Resolution test for SARS-CoV-2 models with *N*_*s*_=26 tetra-rod spikes. The spikes rod length is *l*_*s*_*/R*=0.5 and tetrahedron width is *w*_*s*_*/R*=0.31. The resolution *R/r*_*o*_ and computed reduced coefficients are presented. The resolution adopted in our simulations is highlighted.

### Spikes distribution

We compare the mobility of virions having homogeneously and randomly distributed spikes to elucidate the effects of spikes sparsity. For homogeneous distributions, we localized the spikes isotropically at equidistant positions on the surface of the envelope. For random distribution, we model ten replicas of the virion changing the position of the spikes to obtain characteristic mean mobilities. Even though we are modelling the envelope and spikes as rigidly moving objects, this sampling approach is (up to a good approximation) equivalent to model virions with **S** that diffuses over the surface of **E**. In contrast, isotropically distributed spikes would correspond to **S** with a fixed position and consequently lower conformational freedom. We referred the reader to the SI for a detailed description of the virion construction. As a whole, we identify that virion diffusivity increases when spikes are localized at random positions around the surface (see SI Figure 3. Section 6). For all the spike shapes investigated, we observe an increment on both 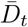 and 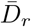. The random distributions of the spikes induce a break in the symmetry of the virion envelope, giving, on average, enhanced mobility. Isotropically distributed spikes preserve the symmetry of the envelopes in a way that the hydrodynamic forces and torques originated due to the presence of spikes is kept balanced across the axes. As the distribution of **S** becomes anisotropic, this balance is broken, favouring virion mobility along some axes. At low spikes densities random configurations exhibit overall lower mobility reduction than homogeneous distributions. As the number of spikes increases, the packing of the spikes become more restricted, and the symmetry-breaking effects of random distributions vanish. Owning the symmetry of spherical envelopes, the effect of spikes distribution appears more significant compared to the ellipsoidal ones.

