## Supporting Information: Method description and calibration. Reported data for various virus. for "Hydrodynamics of spike proteins dictate a transport-affinity competition for SARS-CoV-2 and other enveloped viruses"

#### List of figures and tables

**1. Rigid multiblob methodology.**

**2. Dimensionless diffusivity $D_t$ and $D_r$**

**3. Dimensionless diffusivity $D_t$ and $D_r$ for virions with spherical or ellipsoidal envelope with spikes**

**4. Optimal resolution Determination**

Figure 1: blob radius $ro$ overloaps for sphere and ellipsoids.
Figure 2: Representation of a tetrahedron discretization.

Table 1: Test resolution for a single sphere and ellipsoid.
Table 2: Summary of the favorable resolution for a sphere and ellipsoid.
Table 3: Resolution test of normalized ellipsoid envelope with $N_s$=26 tetra-rod spikes.
Table 4: Different resolution of a discretized tetrahedron.
Table 5: Summary of optimal resolutions of a tetrahedron.

#### 5. Rotational and traslational diffusivity of SARS-CoV-2

Table 6: Comparative translational and rotational diffusivities computed with $D_t$ virion formula and real values of viscosity, temperature and energy for SARS-CoV-2.
Table 7: Actual diffusion of SARS-CoV-2 virion per maximum and minimum spikes number 26 ± 15.

#### 6. Random distribution

Figure 3: Dimensionless rotational and translational diffusivities of random and uniform distribution of the spikes.

Table 8: Fitting results of mean random of ten groups of data, and a single random set of results.

#### 7. Size of the spikes

Figure 4: Dimensionless rotational and translational diffusivity for different spikes length $l_s/R$.
Figure 5: Dimensionless rotational and translational diffusivity for different tetra-spikes radius $R_{\text{tetra}}$.

#### 8. Virions with spikes

Figure 6: Dimensionless rotational diffusivity for the virions.
Figure 7: Dimensionless translational diffusivity for the virions.

Table 9: Morphological parameters of different type of virions with spikes.
Table 10: Spike parameters for the virions according to the type of the spikes and the measurement.
Table 11: Comparative test with different parameters; All spikes distribution is random, the resolution is 14.5, the number and type of spikes are in Table 9.
Table 12: Comparative translational and rotational diffusivities computed with $D_t$ virion formula and real values of viscosity, temperature and energy for all tested viruses.

#### 9. Fitting of the excess diffusivity functions

Figure 8: Schematic of the functional dependence of rotational diffusion coefficient with $N_s$.

Table 13: Fitting constant numbers and Coefficient of determination.

#### 10. Viruses binding affinity information.

Table 14: Binding affinity and $K_d$ for the viruses.

**11. Mesh construction**

Figure 9: Blob representation of spheres using 12, 42, and 162 blobs in a uniform distribution.
Figure 10: Blob representation of tetrahedrons using 10, 34 and 130 blobs in a uniform distribution.
Figure 11: Blob representation of rods using 1, 3 and 7 blobs.

**12. Time scales for virion binding**

Figure 12: Time of transportation in the rotational movement $\tau_r$ in 5e-06 m of mucus.
Figure 13: Time of transportation in the translational movement $\tau_t$ in 5e-06 m of mucus.
Figure 14: Time of transportation ratio $\tau_t/\tau_r$.
Figure 15: Dimensionless rotational diffusivity for different type of spikes.
Figure 16: Dimensionless translational diffusivity for different type of spikes.

### 1 Rigid multiblob methodology

In one of his celebrated papers, Einstein used the linear response theory to demonstrate that the translational diffusion coefficient of a colloid $\Omega$ is related to its translational mobility [1]. The same arguments can be easily extended to include the rotational diffusion [2]. In both cases the diffusion coefficients, $D_t$ and $D_r$, are porportional to the the mobilities,

$$D_t = \frac{k_B T}{3} \mathrm{Tr}\left(\boldsymbol{M}_t\right), \quad D_r = \frac{k_B T}{3} \mathrm{Tr}\left(\boldsymbol{M}_r\right), \tag{1}$$

where $k_B T$ is the thermal energy and Tr denotes the trace operator. The mobility components yield the linear and angular velocities of a colloid ($\boldsymbol{u}$ and $\boldsymbol{\omega}$) in response to applied forces and torques ($\boldsymbol{f}$ and $\boldsymbol{\tau}$),

$$\begin{pmatrix} \boldsymbol{u} \\ \boldsymbol{\omega} \end{pmatrix} = \begin{pmatrix} \boldsymbol{M}_t & \boldsymbol{M}_c \\ \boldsymbol{M}_c^T & \boldsymbol{M}_r \end{pmatrix} \begin{pmatrix} \boldsymbol{f} \\ \boldsymbol{\tau} \end{pmatrix}. \tag{2}$$

For colloids larger than a few nanometers the mobility components can be calculated using the Stokes equations to a good approximation [3, 4]. In this limit, the fluid velocity and pressure, $\boldsymbol{v}$ and $p$, obey the Stokes equations with viscosity $\eta$

$$-\boldsymbol{\nabla} p + \eta \boldsymbol{\nabla}^2 \boldsymbol{v} = 0, \tag{3}$$

$$\boldsymbol{\nabla} \cdot \boldsymbol{v} = 0, \tag{4}$$

while for boundary conditions one can assume that the fluid velocity obeys the no-slip condition at the colloid's surface and that it decays to zero at infinity. We have assumed through this work that the virion, including its spikes, behaves like rigid body, thus the no-slip condition for a virion located at $\boldsymbol{q}$ is quite simple,

$$\boldsymbol{v}(\boldsymbol{r}) = \boldsymbol{u} + \boldsymbol{\omega} \times (\boldsymbol{r} - \boldsymbol{q}) \quad \text{for all } \boldsymbol{r} \in \partial\Omega. \tag{5}$$

These partial differential equations are closed by the balance of force and torque. The integral of the fluid traction, $-\boldsymbol{\lambda}$, over the surface of the virion balance the external forces and torques apply to the virion [5]

$$\int_{\partial\Omega} \boldsymbol{\lambda} \, \mathrm{d}S_r = \boldsymbol{f}, \tag{6}$$

$$\int_{\partial\Omega} (\boldsymbol{r} - \boldsymbol{q}) \times \boldsymbol{\lambda} \, \mathrm{d}S_r = \boldsymbol{\tau}. \tag{7}$$

To solve the Stokes problem and compute the mobilities we use the rigid multiblob method [6]. We discretize the surface of the virion with $N$ markers or *blobs* of radius $a$ and with position $\boldsymbol{r}_i$. The blobs are subject to constraint forces, $\boldsymbol{\lambda}_i$, that ensure the rigid motion of the whole virion. Evaluating the no-slip condition at the blobs, as in collocation methods, leads to a linear system of equations for the unknowns $\boldsymbol{u}$, $\boldsymbol{\omega}$ and $\boldsymbol{\lambda}_i$,

$$\boldsymbol{v}(\boldsymbol{r}_i) = \sum_{j=1}^{N} (\boldsymbol{M}_B)_{ij} \, \boldsymbol{\lambda}_j = \boldsymbol{u} + \boldsymbol{\omega} \times (\boldsymbol{r}_i - \boldsymbol{q}) \quad \text{for } i = 1, \dots, N, \tag{8}$$

$$\sum_{j=1}^{N} \boldsymbol{\lambda}_j = \boldsymbol{f}, \tag{9}$$

$$\sum_{j=1}^{N} (\boldsymbol{r}_j - \boldsymbol{q}) \times \boldsymbol{\lambda}_j = \boldsymbol{\tau}, \tag{10}$$

In the no-slip equation, Eq. (8), the blob mobility matrix $(\boldsymbol{M}_B)_{ij}$ couples the force acting on the blob $j$ to the flow generated at the blob $i$. We use the regularized Rotne-Prager mobility

$$(\boldsymbol{M}_B)_{ij} = \left( \boldsymbol{I} + \frac{a^2}{6} \boldsymbol{\nabla}^2_{\boldsymbol{r}} \right) \left( \boldsymbol{I} + \frac{a^2}{6} \boldsymbol{\nabla}^2_{\boldsymbol{r}'} \right) \boldsymbol{G}(\boldsymbol{r}, \boldsymbol{r}') |_{\boldsymbol{r}' = \boldsymbol{r}_j}^{\boldsymbol{r} = \boldsymbol{r}_i}, \tag{11}$$

where $\boldsymbol{G}(\boldsymbol{r}, \boldsymbol{r}')$ is Green's function of the Stokes equation, i.e. the Oseen kernel. The Rotne-Prager mobility has a closed analytical expression [7, 8]. Given the forces and torques the linear system (8)-(10) can be solved to compute the particle velocities or, equivalently, the mobility matrix components from Eq. (2).

### 2 Dimensionless diffusivity $\bar{D}_t$ and $\bar{D}_r$

The theoretical translational diffusion of a sphere is

$$D_t = \frac{k_B T}{6 \pi r \eta}, \tag{12}$$

and the rotational is

$$D_r = \frac{k_B T}{8\pi r^3 \eta}. \tag{13}$$

Furthermore, the translational mobility is

$$\boldsymbol{M}_t = \frac{1}{6\pi r \eta}, \tag{14}$$

and the rotational mobility is

$$\boldsymbol{M}_r = \frac{1}{8\pi r^3 \eta}. \tag{15}$$

The ratio between computed mobility $\boldsymbol{M}_t$ and theoretical mobility $\boldsymbol{M}_t^o$ is given by,

$$\bar{D}_t = \frac{\boldsymbol{M}_t}{\boldsymbol{M}_t^o} \tag{16}$$

for translational, and

$$\bar{D}_r = \frac{\boldsymbol{M}_r}{\boldsymbol{M}_r^o} \tag{17}$$

for rotational.

### 3 Dimensionless diffusivity $\bar{D}_t$ and $\bar{D}_r$ for virions with spherical or ellipsoidal envelope with spikes

#### 3.1 Sphere with spikes

$\bar{D}_t$ and $\bar{D}_r$ of virions with spikes are calculated as the ratio between the mobility of the virions with spikes $\boldsymbol{M}_t$ for translational and $\boldsymbol{M}_r$ for rotational, divided by the single sphere $\boldsymbol{M}_t|_{\text{sphere}}$ and $\boldsymbol{M}_r|_{\text{sphere}}$ respectively,

$$\bar{D}_t = \frac{\boldsymbol{M}_t}{\boldsymbol{M}_t|_{\text{sphere}}}, \tag{18}$$

$$\bar{D}_r = \frac{\boldsymbol{M}_r}{\boldsymbol{M}_r|_{\text{sphere}}}. \tag{19}$$

The Eq.18 and Eq.19 are the computed dimensionless diffusivities for virions with spherical envelope.

#### 3.2 Ellipsoid with spikes

Similar to the single ellipsoid and the sphere with spikes, in this case, $\bar{D}_t$ and $\bar{D}_r$ are the dimensionless ratio between computed mobility of the particle with spikes $\boldsymbol{M}_t$, $\boldsymbol{M}_r$ divided by the calculated mobility of the single ellipsoid $\boldsymbol{M}_t|_{\text{ellip}}$ and $\boldsymbol{M}_r|_{\text{ellip}}$

$$\bar{D}_t = \frac{\boldsymbol{M}_t}{\boldsymbol{M}_t|_{\text{ellip}}}, \tag{20}$$

and

$$\bar{D}_r = \frac{\boldsymbol{M}_r}{\boldsymbol{M}_r|_{\text{ellip}}}. \tag{21}$$

The Eq.20 and Eq.21 are the computed dimensionless diffusivities for virions with ellipsoid envelope.

### 4 Optimal resolution determination

We calculate an optimal resolution to simulate the virions computing the rotational and translational mobilities of a sphere, ellipsoid, and different spikes morphologies. The optimal resolution is given when the difference between simulations converges. The resolution is given for the radius of the sphere divided by the distance between particles ($ro$)

$$\text{Resolution} = \frac{\text{Radius}}{ro}. \tag{22}$$

We test six different resolutions of 0.9 (12 particles), 1.8 (42 particles), 3.6(162 particles), 7.2(642 particles), 14.5(2562 particles) and 29 (10242 particles).

The errors of $\bar{D}_t$ and $\bar{D}_r$ are

$$\text{Error}_{D_t} = \frac{\boldsymbol{M}_t^o - \boldsymbol{M}_t}{\boldsymbol{M}_t}$$

and

$$\text{Error}_{D_r} = \frac{\boldsymbol{M}_r^o - \boldsymbol{M}_r}{\boldsymbol{M}_r},$$

in Table 1. we registered the error for all the resolutions, and choose the favorable resolution as the best trade off between accuracy and computational cost. The resolution of 29 provided the better approximation, with a rather high computational cost. Therefore, we selected as a reasonable resolution the corresponding 14.5, with errors on the order of $1\%$ in $\bar{D}_t$, and $3\%$ for $\bar{D}_r$, for an spherical envelope. We remark that acceptable results can be already obtained with resolutions of 7.2, having errors of $2\%$ for $\bar{D}_t$, and $6\%$ for $\bar{D}_r$.

#### 4.1 Ellipsoid - Equivalent radius $R_e$

For ellipsoids, we calculated an equivalent radius $R_e$ of a sphere with equivalent volume. The volume of an ellipsoid with principal axis length $a$, $b$, and $c$, is

$$V_{\text{ellip}} = \frac{4}{3}\pi abc. \tag{23}$$

The equivalent radius is then given by

$$R_e = \left(\frac{3}{4\pi}V_{\text{ellip}}\right)^{\frac{1}{3}}. \tag{24}$$

##### 4.2 Optimal resolution test

To determine an optimal resolution, are tested six different resolutions for sphere and ellipsoid envelopes. To calculate the mobility is important to consider the blob radius for the spheres and ellipsoids (section 4.2.1).

Table 1: Test resolution for a single sphere and ellipsoid

| **Resolution** | **Number of particles** | **Envelope** | $M_t$ | $M_r$ | $R_e$ | $\frac{M_t^o - M_t}{M_t}$ | $\frac{M_r^o - M_r}{M_r}$ | **Figure** |
|---|---|---|---|---|---|---|---|---|
| 29 | 10242 | Sphere | 0.0527 | 0.0391 | - | 0.005 | 0.016 | - |
| 14.5 | 2562 | Sphere | 0.0524 | 0.0384 | - | 0.012 | 0.035 | 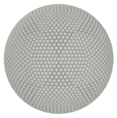 |
| 7.2 | 642 | Sphere | 0.0518 | 0.0372 | - | 0.024 | 0.065 | 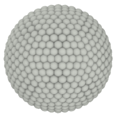 |
| 3.6 | 162 | Sphere | 0.0503 | 0.0304 | - | 0.052 | 0.234 | 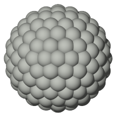 |
| 1.8 | 42 | Sphere | 0.0472 | 0.0297 | - | 0.110 | 0.254 | 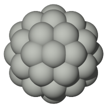 |
| 0.9 | 12 | Sphere | 0.0420 | 0.0213 | - | 0.208 | 0.465 | 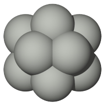 |
| 29 | 10242 | Ellipsoid | 0.0609 | 0.0595 | 0.8671 | 0.004 | 0.025 | - |
| 14.5 | 2562 | Ellipsoid | 0.0600 | 0.0570 | 0.8770 | 0.008 | 0.033 | 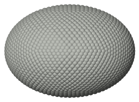 |
| 7.2 | 642 | Ellipsoid | 0.0585 | 0.0534 | 0.8967 | 0.012 | 0.033 | 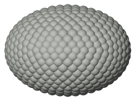 |
| 3.6 | 162 | Ellipsoid | 0.0550 | 0.0460 | 0.9360 | 0.030 | 0.051 | 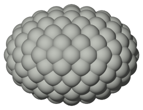 |
| 1.8 | 42 | Ellipsoid | 0.0476 | 0.0323 | 1.0134 | 0.091 | 0.156 | 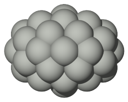 |
| 0.9 | 12 | Ellipsoid | 0.0392 | 0.0152 | 1.157 | 0.144 | 0.406 | 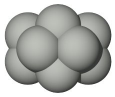 |

Table 2: Summary of the favorable resolution for a sphere and ellipsoid

| Test | Envelope | Resolution | $R_x$ | $R_y$ | $R_z$ | $R_t$ | $R_r$ | $\boldsymbol{M}_t$ | $\frac{\boldsymbol{M}_t^o - \boldsymbol{M}_t}{\boldsymbol{M}_t}$ | $\boldsymbol{M}_r$ | $\frac{\boldsymbol{M}_r^o - \boldsymbol{M}_r}{\boldsymbol{M}_r}$ |
|---|---|---|---|---|---|---|---|---|---|---|---|
| **1** | Sphere | 14.5 | 1 | 1 | 1 | 1.057 | 1.050 | 0.052 | 0.012 | 0.038 | 0.035 |
| **2** | Ellipsoid | 14.5 | 1 | 0.9 | 0.7 | 0.927 | 0.923 | 0.060 | 0.008 | 0.057 | 0.033 |

###### 4.2.1 Blob radius

In the computational methodology RMB, we consider particles are located at a distance $r_o$, and set a blob radius of $r_b = ro/2$ for spheres and $r_b = 0,75ro$ for ellipsoids to ensure the accuracy of the method.

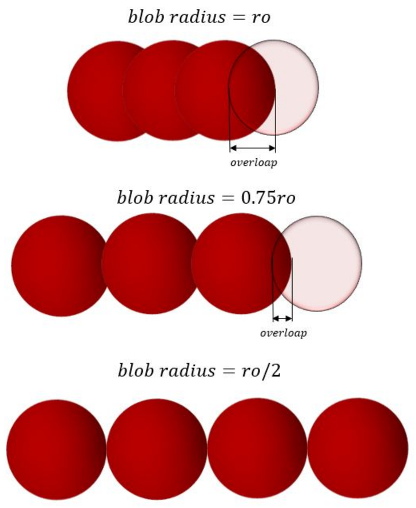


Figure 1: blob radius $ro$ have overloaps, to minimize the overloap is defined a blob radius of $ro/2$ for sphere, and $0.75ro$ for ellipsoids.

##### 4.3 Resolution envelope and spikes for SARS-CoV-2

We test four different resolution using spikes and envelope with the SARS-CoV-2 configuration. The main is identify the resolution differences using the spikes and envelope together. In table 3, is illustrated the outcomes for the dimensionless diffusivity and the computed radius. Considering the resolution of 29 as the best approximation, an optimal resolution of 14.5 have a maximum error of $0.7\%$. This is in contrast with coarser resolutions of 3.6 with errors on the order of $20\%$ for $\bar{D}_r$.

Table 3: Resolution test of normalized ellipsoid envelope with $N_s$=26 tetra-rod spikes. The spikes length is $l_s/R$=0.5 and tetrahedron width is $w_s/R$=0.31. Where $R$ is the largest axis of the ellipsoidal envelope.

| **Resolution** | $\bar{D}_t$ | $\bar{D}_r$ | $R_t$ | $R_r$ | **Figure** |
|---|---|---|---|---|---|
| 29 | 0.7682 | 0.4162 | 1.133 | 1.171 | 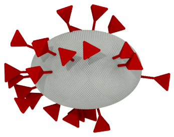 |
| 14.5 | 0.7638 | 0.4131 | 1.157 | 1.190 | 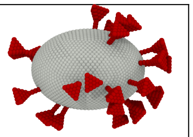 |
| 7.2 | 0.7478 | 0.3882 | 1.213 | 1.243 | 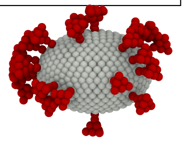 |
| 3.6 | 0.7075 | 0.3303 | 1.364 | 1.378 | 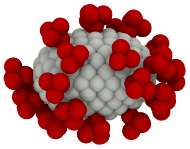 |

#### 4.4 Tetrahedral spike resolution

For tetrahedral spikes we also conduct a separate calibration of the tetrahedron alone. We consider a tetrahedron of with $a = w_s$, inter-blob distance $r_o$, height is $H_{\text{tetra}} = \frac{\sqrt{6}}{3}a$, and circumscribed in a sphere of radius $R_{\text{tetra}} = w_s(3/8)^{1/2}$ See Figure 2. Then, we characterize them using its equivalent radius ($R_e$), defined by the radius of a sphere with equivalent volume of the tetrahedron, $V_{tetra} = \frac{a^3}{6\sqrt{2}}$, leading to

$$R_e = \left(\frac{3}{4\pi}V_{\text{tetra}}\right)^{\frac{1}{3}}. \quad (25)$$

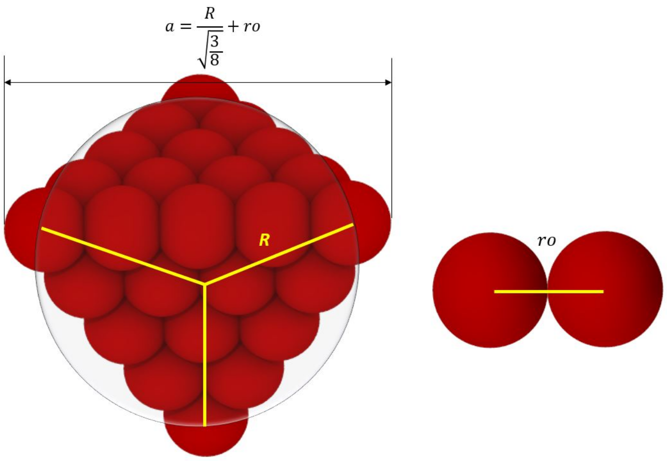


Figure 2: Representation of a tetrahedron of width $a$, inter-blob distance $r_o$, and circumscribed radius $R_{\text{tetra}} = w_s(3/8)^{1/2}$.

The diffusivities for the single spikes are reduced as for the spheres and ellipsoids, with respect to a reference mobility $\boldsymbol{M}_t^o$, such that

$$\bar{D}_t^{tetra} = \frac{\boldsymbol{M}_t^{\text{tetra}}}{\boldsymbol{M}_t^o},$$

$$\bar{D}_r^{tetra} = \frac{\boldsymbol{M}_r^{\text{tetra}}}{\boldsymbol{M}_r^o}.$$

Since we do not count with an analytical expression for the mobility of a tetrahedron, we define the reference mobility, as the one of a sphere of radius $R_e|_{\text{solid}}$, where $R_e|_{\text{solid}}$ is the equivalent radius of a solid tetrahedron of width $a$. In table 4, we present the computed mobilities for different resolutions $R_{\text{tetra}}/r_o$, and the convergence criteria for each case. As a convergence criteria, we use the relative difference with respect to the highest resolution achieved, before the calculation become computationally prohibitive. Using a spike resolution of 2.5 we obtain an error of $3\%$ for $M_t$ and $8\%$ for $M_r$, further improvement is achieved with resolutions of 4.9 with $M_t$ and $M_r$ errors of $1\%$ and $4\%$ respectively. The better resolutions reduce the error, but, requires a higher computational cost.

Table 4: Different resolution of a discretized tetrahedron. The mobilities for the different resolutions are reduced using the mobility of a sphere of radius $R_e$, for a solid tetrahedron of edge $a$. As a convergence estimator, we measure variation on the reduced mobility with respect to the largest resolution simulated.

| Resolution | Number of particles | $\boldsymbol{M}_t/\boldsymbol{M}_t^o$ | $\boldsymbol{M}_r/\boldsymbol{M}_r^o$ | $R_e$ | $a$ | $\frac{\boldsymbol{M}_{t\max}-\boldsymbol{M}_t}{\boldsymbol{M}_{t\max}}$ | $\frac{\boldsymbol{M}_{r\max}-\boldsymbol{M}_r}{\boldsymbol{M}_{r\max}}$ | Figure |
|---|---|---|---|---|---|---|---|---|
| Solid tetrahedron | | – | – | 0.4966 | 1.6330 | 0 | 0 | |
| 40 | 8194 | 0.8215 | 0.4750 | 0.5042 | 1.6580 | 0 | 0 | |
| 19.6 | 2050 | 0.8200 | 0.4724 | 0.5121 | 1.6840 | 0.0019 | 0.0055 | 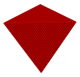 |
| 9.8 | 514 | 0.8159 | 0.4660 | 0.5276 | 1.7350 | 0.0069 | 0.0191 | 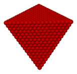 |
| 4.9 | 130 | 0.8074 | 0.4541 | 0.5587 | 1.837 | 0.0172 | 0.0440 | 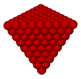 |
| 2.5 | 34 | 0.7918 | 0.4370 | 0.6207 | 2.0410 | 0.0362 | 0.0801 | 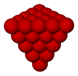 |
| 1.2 | 10 | 0.7651 | 0.4235 | 0.7448 | 2.4490 | 0.0688 | 0.1068 | 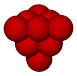 |

Table 5: Summary of optimal resolutions of a tetrahedron. Low resolutions of 4.9 and 2.5 are the best options because of the computational cost of the simulations.

| Resolution | Number of particles | Error$_{M_t}$ | Error$_{M_r}$ |
|---|---|---|---|
| 4.9 | 130 | 1.7 | 4.4 |
| 2.5 | 34 | 3.6 | 8.0 |

#### 5 Rotational and traslational diffusivity of SARS-CoV-2

For practical applications, virion diffusivities, $D_t|_{\text{virion}}$, can be easily retrieved using the known diffusivities of their respective plain envelope (sphere or ellipsoid), $D_i|_{\text{envelope}}$, and the reduced diffusivity $\bar{D}_t$, such that

$$D_t|_{\text{virion}} = \bar{D}_t D_t|_{\text{envelope}}, \tag{26}$$

$$D_t|_{\text{envelope}} = \frac{k_B T}{6\pi\eta R_{\text{env}}}. \tag{27}$$

For SARS-CoV-2, we compute the diffusivities for G-form and D-form, the results are in Table 6.

Table 6: Comparative translational and rotational diffusivities computed with $D_t^{\text{virion}}$ formula and real values of viscosity, temperature and energy. The viscosity values are for water = 0.001 Pa · s, blood = 0.0035 Pa · s [9] and, nasal mucus = 1.6 Pa · s [10]. Temperature = 298.15 K, and kb = 1.38064852e-23 $\text{m}^2\text{kg}/\text{s}^2\text{K}$

| | water | | blood | | nasal mucus | |
|---|---|---|---|---|---|---|
| Virus | $D_t$ $[\mu\text{m}^2/\text{s}]$ | $D_r$ $[1/\text{s}]$ | $D_t$ $[\mu\text{m}^2/\text{s}]$ | $D_r$ $[1/\text{s}]$ | $D_t$ $[\mu\text{m}^2/\text{s}]$ | $D_r$ $[1/\text{s}]$ |
| SARS-CoV-2 (G-form) | 4.0451 | 965.4694 | 1.1557 | 275.8484 | 2.5281 e-3 | 0.6034 |
| SARS-CoV-2 (D-form) | 4.3345 | 1185.2682 | 1.2414 | 338.6481 | 2.7156 e-3 | 0.7408 |

Since the range in the number of spikes reported for SARS-CoV-2 is around $26 \pm 15$, for completness, we compute the diffusivities variation over that range as presented in Table 7.

Table 7: Actual diffusion of SARS-CoV-2 virion per maximum and minimum spikes number $26 \pm 15$

| $N_s$ | $D_t$ $[\mu^2/\text{s}]$ | $D_r$ $[1/\text{s}]$ |
|---|---|---|
| 11 | 4.4842 | 1703.2178 |
| 26 | 4.0451 | 965.4694 |
| 41 | 4.0132 | 953.7624 |

### 6 Random distribution

The random distribution of spikes brings different mobility results. We compute ten replicas using the SARS-CoV-2 morphology with different number of spikes (12,15,18,21,24,26,30,33,36,39,42,50 and 100) and a resolution of 14.5, to estimate the error bars with the standard deviation. We calculate the fitting curve using the function in the Eq 28 of the mean of random replicas and just one of the replicas. The difference between the fitting functions illustrates an error of 2% in the number of spikes.

Table 8: Fitting results of mean random of ten groups of data, and a single random set of results. The error is of 2% in the maximum number of spikes

| **Mobility** | **TRANSLATIONAL** | | | **ROTATIONAL** | | |
|---|---|---|---|---|---|---|
| | $N_s^\infty$ | b | $\bar{D}_t^\infty$ | $N_s^\infty$ | b | $\bar{D}_r^\infty$ |
| Mean random | 141,6 | 10,7 | 0,670 | 105,2 | 6,1 | 0,304 |
| Random | 144,4 | 10,9 | 0,670 | 107,1 | 6,2 | 0,303 |
| Error | 1,9 | 1,8 | 0,000 | 1,7 | 2,5 | 0,330 |

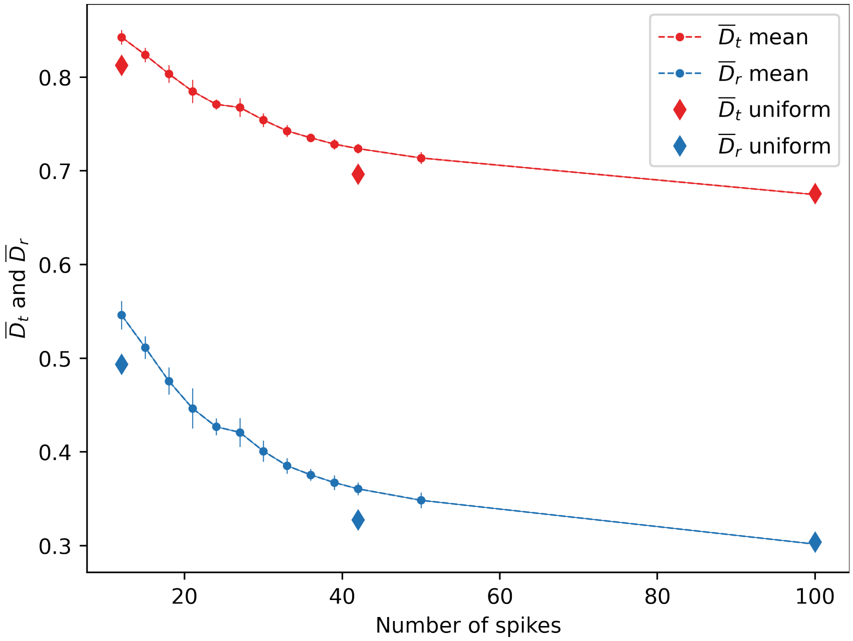


0

Figure 3: Dimensionless rotational and translational diffusivities of random and uniform distribution of the spikes. For random distribution we obtain the mean and the error bars for 12,15,18,21,24,26,30,33,36,39,42,50 and 100 spikes. On the other hand, for uniform distribution we obtain diffusivities for 12, 42 and 162 spikes because the uniformity restrict the number of spikes. The values of 100 spikes is calculated with an interpolation between 42 and 162 values. However, it is evident that at large $N_s$, both random and uniform distribution converge due to the packing of the spikes in the envelope.

In Figure 3 we present the dimensionless diffusivities of a SARS-CoV-2, using random and homogeneous distributions with different number of spikes. For the homogeneous distributions we required equidistant location of the spikes, thus we tested $N_s$ equal to 12, 42, and 162. Models with 100 spikes cannot be constructed directly without further optimization on spikes location, therefore for simplicity we calculated it as the interpolation between $N_s = 42$ and $N_s = 162$. For the random distribution are illustrated the mean of the diffusivities and the error bars. We can observe the error bars reduced as the number of spikes increases. Furthermore, the homogeneous distribution leads to diffusivities that are approximately $9.6\%$ smaller than the random ones. However, as the number of spikes increases, the diffusivities of both homogeneous and random converged (difference of around $0.74\%$) due to the crowding of the spikes, leading to a close packing.

### 7 Size of the spikes

If the length and width of the spikes increase, $\bar{D}_t$ and $\bar{D}_r$ diminishes in a consistent fashion. Moreover, the rotational diffusivity exhibits a greater reduction than the translational one because of its dependency on the radius. Both, the length and the width are dimensionless, divided by the envelope radius. In Figure 4 and 5, we present the variation on both $\bar{D}_t$ and $\bar{D}_r$ for virion models with rod and tetra spikes, respectively. For rod type the size $l_s$ is varied, whereas for tetra type the radius $R_{\text{tetra}}$ (see Figure 2) is varied. We can observe that the dimensionality of the spikes affects the scaling of diffusion with the spike size. Comparing Figure 4 and 5, rod shapes (that are dominantly one dimensional) showed a weaker variation on $\bar{D}_t$ and $\bar{D}_r$ as the size increases. In contrast, tetra-shape spikes displayed a strong reduction on diffusivity. Overall, we observed that the virions with bulkier and larger spikes (compared to the envelope size) had an intrinsic diffusional penalty. Therefore, the regulation on the number of spikes suggests a possible alternative to compensate for this reduction in mobility.

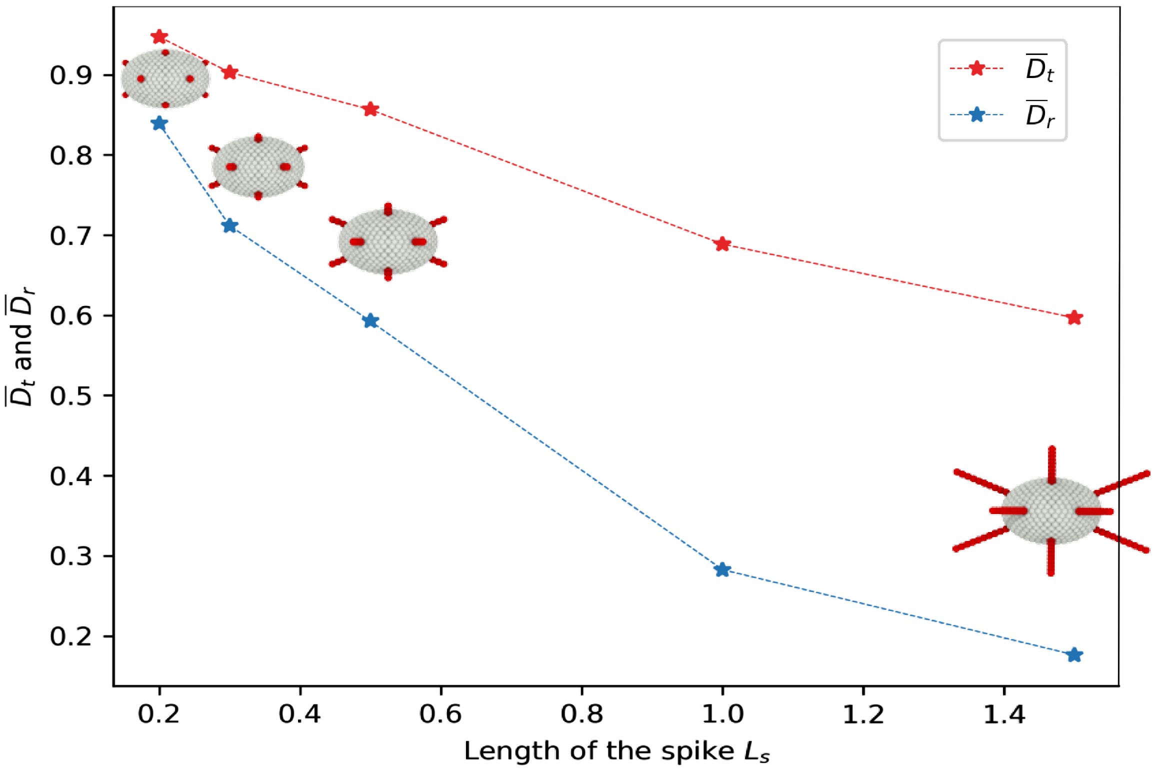


Figure 4: Dimensionless rotational and translational diffusivity for different spikes length $l_s/R$. The simulation is with ellipsoid envelope, $N_s$=12, uniform distribution and rod spikes.

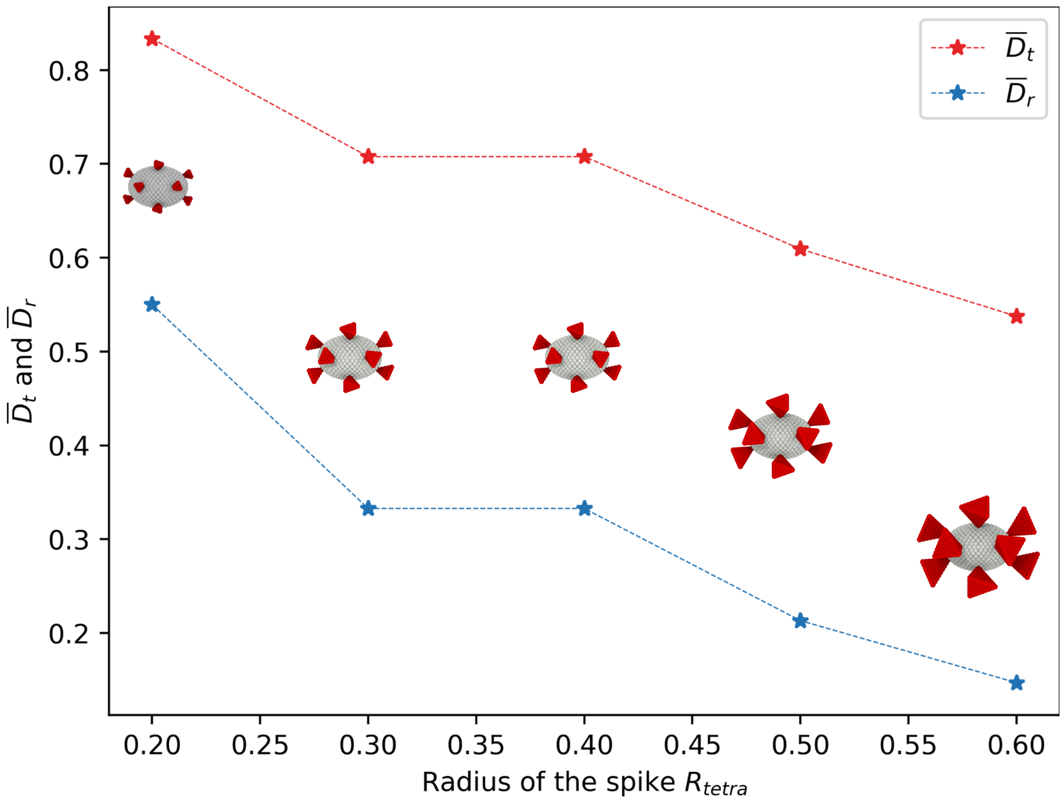


Figure 5: Dimensionless rotational and translational diffusivity for different tetra-spikes radius $R_{\text{tetra}}$. The simulation is with ellipsoid envelope, $N_s$=12, uniform distribution and tetrahedron spikes.

#### 8 Virions with spikes

Other viruses with similar morphologies as SARS-CoV-2 are tested and compared. The tested virions are SARS-CoV-2, HIV, MHV, SARS-CoV, Dengue, Lassa, Influenza and Herpes Simplex.

Table 9: Morphological parameters of different type of virions with spikes

| Virus | Radius envelope [nm] | Number of spikes | Length of spikes [nm] | Width of spikes [nm] | Type of spike | Reference |
|---|---|---|---|---|---|---|
| SARS-CoV2 | 48 | 26 | 25 | 15 | Tetra rod | [11] |
| HIV | 60 | 14 | 12 | 15 | Tetra | [12] |
| Dengue | 20 | 60 | 11 | 4 | Tetra rod | [13] |
| Lassa | 66 | 273 | 9 | 10 | Sphere rod | [14] |
| Influenza | 60 | 350 | 15 | 7 | Rod | [15] |
| Herpes Simplex | 93 | 659 | 20 | 5 | Rod | [16] |
| SARS-CoV | 50 | 65 | 16 | 18 | Tetra | [17] |
| MHV | 42.5 | 11 | 20 | 10 | Sphere rod | [18] |

Table 10: Spike parameters for the virions according to the type of the spikes and the measurement. The dimensionless length and width are normalized with the radius of the virion, and are used to calculate the approximated total length of the spike $S\_s = (L\_s/R + W\_s/R + ro)$. $S\_s$ is used in the simulations.

| Virions | Type of spike | Dimensionless Length $L/R$ | Dimensionless Width $W/R$ | $S\_s$ |
|---|---|---|---|---|
| SARS-CoV-2 | Tetra rod | 0,52 | 0,31 | 0,59 |
| HIV | Tetra | 0,20 | 0,25 | 0,34 |
| Dengue | Tetra rod | 0,55 | 0,20 | 0,57 |
| Lassa | Sphere rod | 0,14 | 0,15 | 0,34 |
| Influenza | Rods | 0,25 | 0,12 | 0,32 |
| Herpes Simplex | Rods | 0,22 | 0,05 | 0,27 |
| SARS-CoV | Tetra | 0,32 | 0,36 | 0,36 |
| MHV | Sphere rod | 0,47 | 0,24 | 0,51 |

Table 11: Comparative test with different parameters. All spikes distribution is random, the resolution is 14.5, the number and type of spikes are in Table 9.

| **Test** | **Envelope** | $R_x$ [nm] | $R_y$ [nm] | $R_z$ [nm] | $R_t$ [nm] | $R_r$ [nm] | $\bar{D}_t$ | $\bar{D}_r$ | Figure |
|---|---|---|---|---|---|---|---|---|---|
| **SARS-CoV-2**[11] | Ellips. | 48 | 43 | 33 | 55,675 | 57,275 | 0,762 | 0,410 | 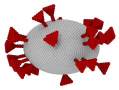 |
| **HIV**[12] | Sphere | 60 | 60 | 60 | 65,830 | 66,539 | 0,922 | 0,759 | 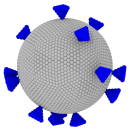 |
| **Dengue**[13] | Sphere | 20 | 20 | 20 | 27,039 | 27,532 | 0,749 | 0.397 | 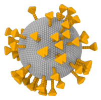 |
| **Lassa** [14] | Sphere | 66 | 66 | 66 | 78,227 | 78,442 | 0,854 | 0,617 | 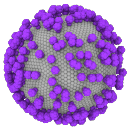 |
| **Influenza** [15] | Sphere | 60 | 60 | 60 | 70,253 | 70,472 | 0,864 | 0,6394 | 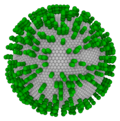 |
| **Herpes S.** [16] | Sphere | 93 | 93 | 93 | 105.000 | 105.129 | 0.896 | 0.717 | 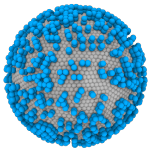 |
| **SARS-CoV** [17] | Sphere | 50 | 50 | 50 | 61.996 | 53.569 | 0.816 | 0.533 | 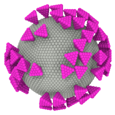 |
| **MHV** [18] | Sphere | 42.5 | 42.5 | 42.5 | 53.040 | 46.075 | 0.944 | 0.813 | 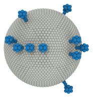 |

Diffusivities per number of spikes are illustrated in the figures 6 for rotational and figure 7 for translational.

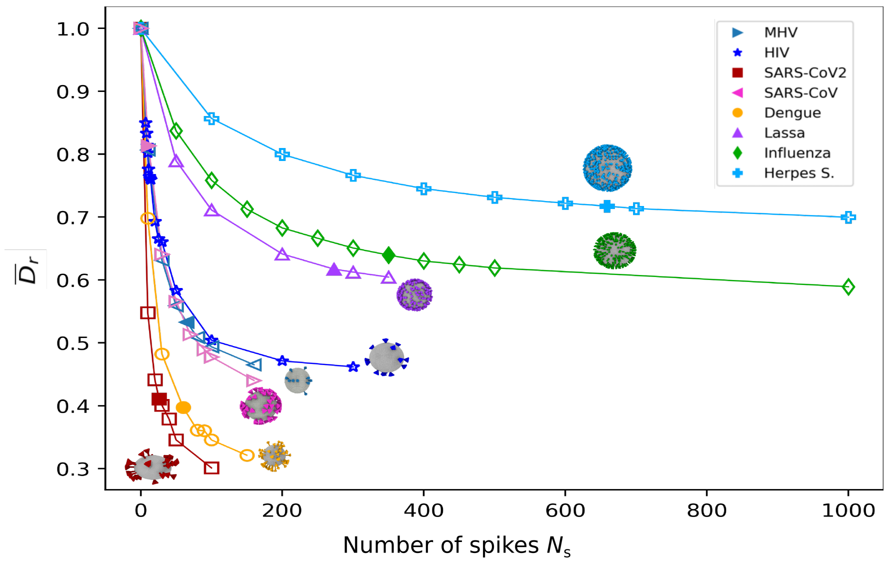


Figure 6: Dimensionless rotational diffusivity for the virions. The coloured marker is the number of the spikes of the virion.

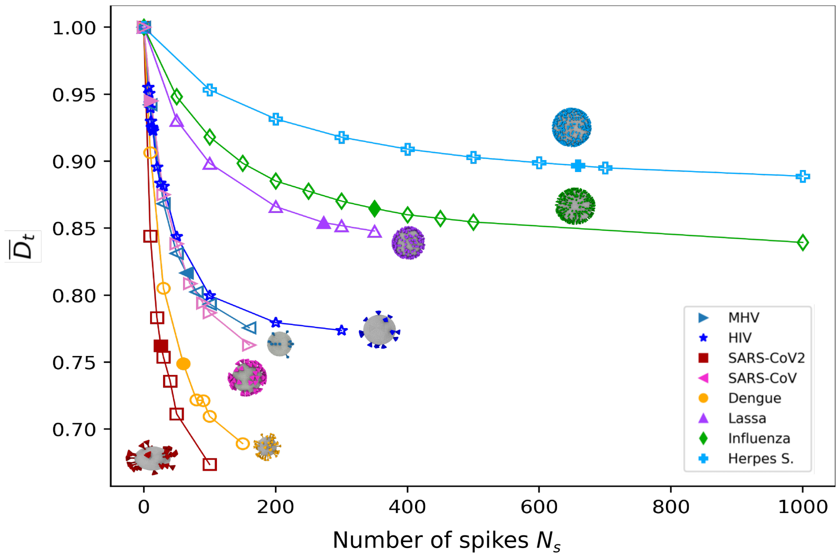


Figure 7: Dimensionless translational diffusivity for the virions. The coloured marker is the number of the spikes of the virion

Table 12: Comparative translational and rotational diffusivities computed with $D_t^{\text{virion}}$ formula and real values of viscosity, temperature and energy. The viscosity values are for water = 0.001 Pa · s, blood = 0.0035 Pa · s [9] and, nasal mucus = 1.6 Pa · s [10]. Temperature = 298.15 K, and kb = 1.38064852e-23 $\mathrm{m^2 kg/s^2 K}$

| | water | | blood | | nasal mucus | |
|---|---|---|---|---|---|---|
| Virus | $D_t$ $[\mu\mathrm{m}^2/\mathrm{s}]$ | $D_r$ $[1/\mathrm{s}]$ | $D_t$ $[\mu\mathrm{m}^2/\mathrm{s}]$ | $D_r$ $[1/\mathrm{s}]$ | $D_t$ $[\mu\mathrm{m}^2/\mathrm{s}]$ | $D_r$ $[1/\mathrm{s}]$ |
| SARS-CoV2 | 4.0451 | 965.4694 | 1.1557 | 275.8484 | 2.5281 e-3 | 0.6034 |
| HIV | 3.3594 | 576.2858 | - | - | - | - |
| Dengue | 8.1784 | 8127.9047 | - | - | - | - |
| Lassa | 2.8257 | 351.5049 | - | - | - | - |
| Influenza | 3.1483 | 484.5350 | - | - | - | - |
| Herpes Simplex | 2.1063 | 145.9985 | - | - | - | - |
| SARS-CoV | 3.852 | 843.828 | - | - | - | - |
| MHV | 4.532 | 1374.032 | - | - | - | - |

### 9 Fitting of the excess diffusivity functions

The results were plotted according the normalized diffusion $\bar{D}_t$ and $\bar{D}_r$. For example:

$$\bar{D}_i = \bar{D}_i^{\infty}(1 - \frac{(N_s^{\infty} - x)}{N_s^{\infty}}(1 - e^{\frac{-b}{x}})) + \frac{N_s^{\infty} - x}{N_s^{\infty}}(1 - e^{\frac{-b}{x}}), \tag{28}$$

where $N_s^{\infty}$ is the number of spikes at which the virion mobility saturates, b is a fitting parameter that depend on the characteristic size of the spike and, $\bar{D}_i^{\infty}$ is the diffusivity of the virus with me maximum number of spikes in the model simulation (Figure 8).

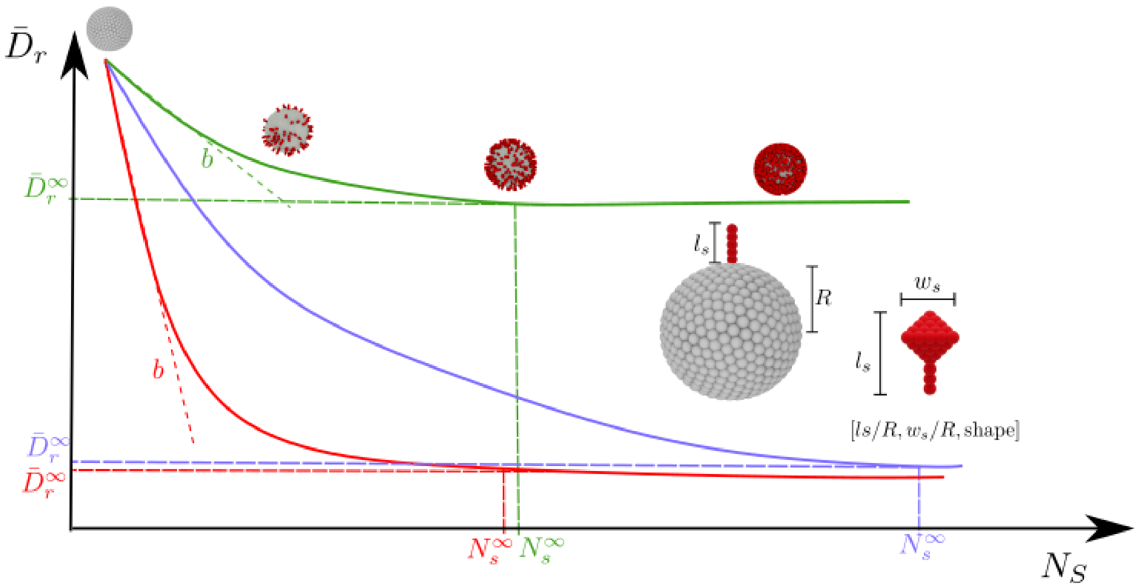


Figure 8: Schematic of the functional dependence of rotational diffusion coefficient with $N_s$. The shape-dependent parameters $N_s^\infty$ and $b$ characterize the behavior of the diffusional decay. $N_s^\infty$ indicates the saturation in mobility when the diffusion is no longer affected by $N_s$, whereas $b$ sets the strength on the decay. Lower values of $b$ correlates with bulkier spikes that induce a fast reduction in the mobility. Moreover the magnitude of $b$ can be used as a differentiating parameter for virion classification.

To determine the accuracy of the proposed fitting equation we computed the coefficient of determination $R_d^2$. The mean $\bar{y}$ is

$$\bar{y} = \frac{1}{n}\sum_{i=1}^{n} y_i, \tag{29}$$

and the coefficient is

$$R_d^2 = 1 - \frac{\sum(y_i - f_i)^2}{\sum(y_i - \bar{y})^2}. \tag{30}$$

According the Eq.30 the calculated data $f_i$ is the fitting function (Eq. 28), the observed data $y_i$ are the dimensionless values of diffusion ($\bar{D}_i$), and the mean of the observed data $\bar{y}$ (Eq. 29) is for the dimensionless diffusion. The results summarized in Table 13 show values of $R_d = 0.99$ between dimensionless diffusion and the fitting function introduced.

Table 13: Fitting constant numbers Eq 28 $N_s^\infty$, b, $\bar{D}_i^\infty$, and $R^2$ Coefficient of determination

| | | TRANSLATIONAL | | | | ROTATIONAL | | | |
|---|---|---|---|---|---|---|---|---|---|
| Virus | $N_s$ | $N_s^\infty \pm 2$ | b $\pm 2$ | $\bar{D}_t^\infty$ | $R_d^2$ | $N_s^\infty \pm 2$ | b $\pm 3$ | $\bar{D}_r^\infty$ | $R_d^2$ |
| SARS-CoV2 | 26 [11] | 204.6 | 8.9 | 0.673 | 0.99 | 206.5 | 4.9 | 0.301 | 0.99 |
| HIV | 14 [12] | 110.4 | 15.9 | 0.799 | 0.99 | 144.5 | 10.8 | 0.499 | 0.99 |
| Dengue | 60 [13] | 229.3 | 14.7 | 0.689 | 0.99 | 196.9 | 9.1 | 0.321 | 0.99 |
| Lassa | 273 [14] | 374.4 | 53.2 | 0.847 | 0.99 | 372.7 | 40.9 | 0.604 | 0.99 |
| Influenza | 350 [15] | 579.3 | 70.3 | 0.854 | 0.99 | 580.9 | 54.1 | 0.619 | 0.99 |
| Herpes Simplex | 659 [16] | 753.4 | 115.6 | 0.895 | 0.99 | 843.8 | 93.8 | 0.710 | 0.99 |
| SARS-CoV | 65 [17] | 169.0 | 17.9 | 0.770 | 0.99 | 211.1 | 11.8 | 0.469 | 0.99 |
| MHV | 11 [18] | 157.7 | 23.6 | 0.769 | 0.99 | 171.1 | 14.8 | 0.449 | 0.99 |

### 10 Viruses binding affinity information.

#### 10.1 Spikes concentration

We approximate the molar concentration of spikes as $[S]$ =moles $\mathbf{S}/l$, to compute the saturation function $[S]/([S] + K_D)$ using the number of spikes $N_s$, the length of the spike $l_s$ and the radius of the envelope $R$. The reaction volume is given by $4/3\pi((R + l_S)^3 - R^3)$, and the number of moles $N_s$/NA. In SI.Table 14, we summarize the magnitude of the binding affinities between spikes proteins and cell receptors for the different viruses modelled. Since the smaller the value of $K_D$, the smaller the amount of spikes needed to achieve an effective binding. Viruses with lower $K_D$ will reach saturation earlier.

Table 14: Binding affinity and $K_d$ for the viruses

| **Virus** | **Biding affinity** | $K_d$ [M] | **Reference** |
|---|---|---|---|
| SARS-CoV-2 | ACE-2 | 44e-9 | [19, 20] |
| HIV | CD4 with gp120 | 1.02e-9 | [21] |
| Dengue | CLEC5A | 172e-6 | [22] |
| Lassa | GPC42-50,GPC60-68, and GPC441-449, | 98e-9 | [23] |
| Influenza | Ha-SA | 10e-5 | [24] |
| Herpes Simplex | gD and nectin-1 | 54e-9 | [25] |
| SARS-Cov | ACE2 | 150e-9 to 300e-9 | [26] |
| MHV | NTD | 21.4 e-9 | [27] |

### 11 Mesh construction

#### 11.1 Sphere

The construction of a sphere of radius $R$ and distance between blobs $r_b$ starts with the coarser representation of an icosphere with 12 vertex located at $R$ and distance $r_o$ between them, as shown in Figure SI.9. One step refinement of this structure is conducted by taking the middle points of the segments connecting two adjacent vertex, and projecting those points radially, such that their new position satisfies $R^2 = x^2 + y^2 + z^2$. This refining procedure can be repeated until the distance between vertex is smaller or equal to the target inter-particle distance ($r_o \leq r_b$).

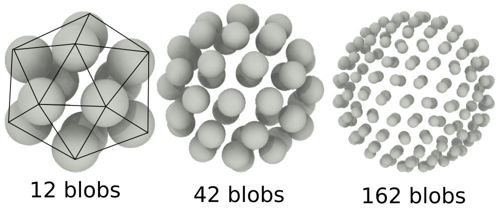


Figure 9: Blob representation of spheres using 12, 42, and 162 blobs in a uniform distribution

#### 11.2 Tetrahedron

The tetrahedron construction starts with the primary coarse surface with only four vertex. Following the same methodology that for the spheres, the structure is refined by splitting in half the edges between two vertex and adding a new point in that position. This addition must be applied to all the edges of the primary surface. This procedure is repeated until the distance between adjacent points is smaller or equal that the target resolution. Tetrahedron with different three different resolutions are presented in SI.10

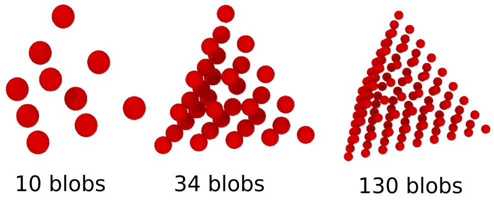


Figure 10: Blob representation of tetrahedrons using 10, 34 and 130 blobs in a uniform distribution.

#### 11.3 Rods

The rods construction consist in linking beads one by one, equidistant and all the centroids in the same angle.

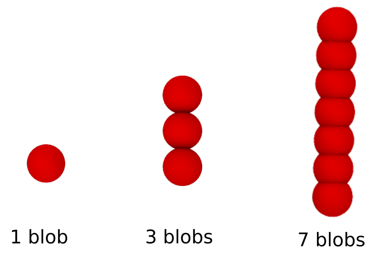


Figure 11: Blob representation of rods using 1, 3 and 7 blobs

#### 12 Time scales for virion binding

We compute the time of transportation of the virions in 5e-06 m of mucus, according the rotational and translational diffusivities. The results illustrated in Figure 12 for rotational, and Figure 13 for translational, show a dependency of the radius of the virion. For example, the biggest virion is the Herpes simplex, and the times ($\tau_t$) and ($\tau_r$) are the largest in contrast to the other virions transportation time.

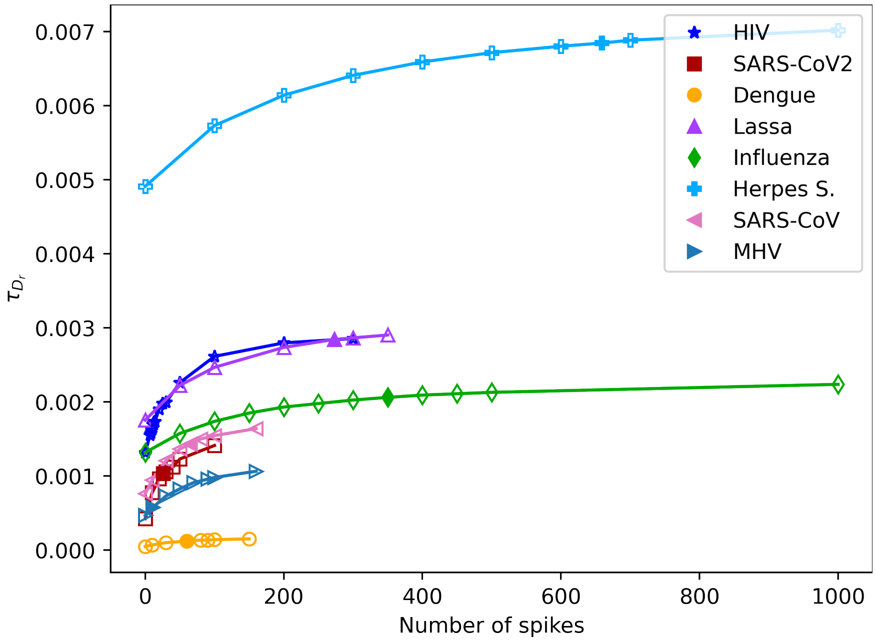


Figure 12: Time of transportation in the rotational movement $\tau_r$ in 5e-06 m of mucus

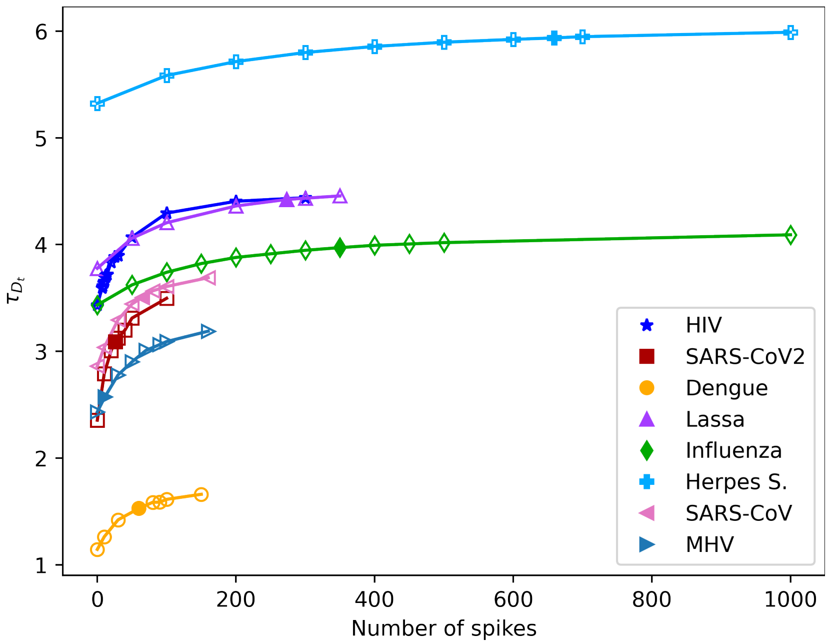


Figure 13: Time of transportation in the translational movement $\tau_t$ in 5e-06 m of mucus

In figure 14 we have the ratio between the transportations time $\tau_t$ and $\tau_r$, the results show a scale of 1000 seconds for the majority of the virions, but for the herpes simplex the scale is in 100 seconds and for dengue is for 10000 seconds. These differences are given by the radius of the virion.

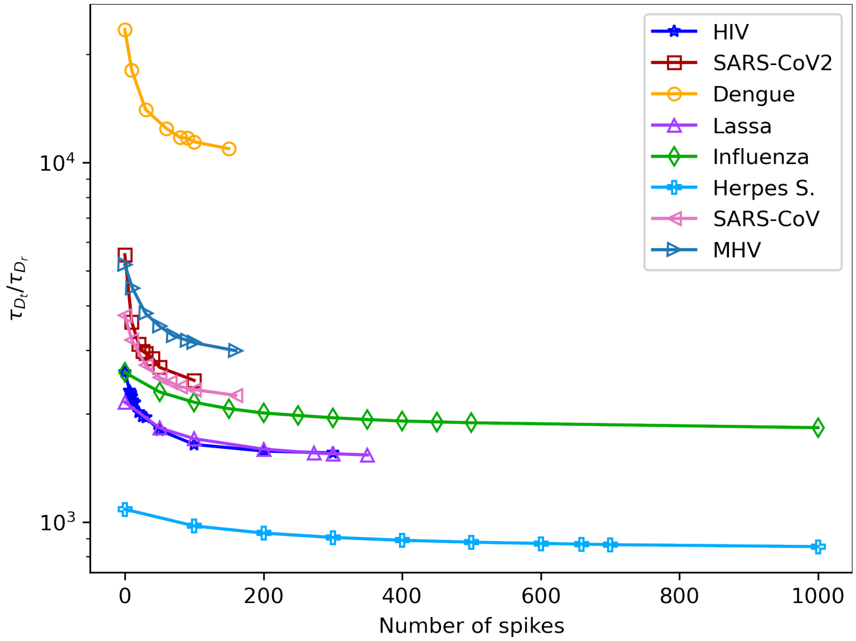


Figure 14: Time of transportation ratio $\tau_t/\tau_r$

### 13 Type of spikes

Five types of spike morphologies are constructed using the RMB methodology: rod, tetra, sphere, rod-tetra, and rod-sphere. Rod, tetra, and sphere are characterize by a single lenght, whereas rod-tetra and rod-sphere require two. parameters

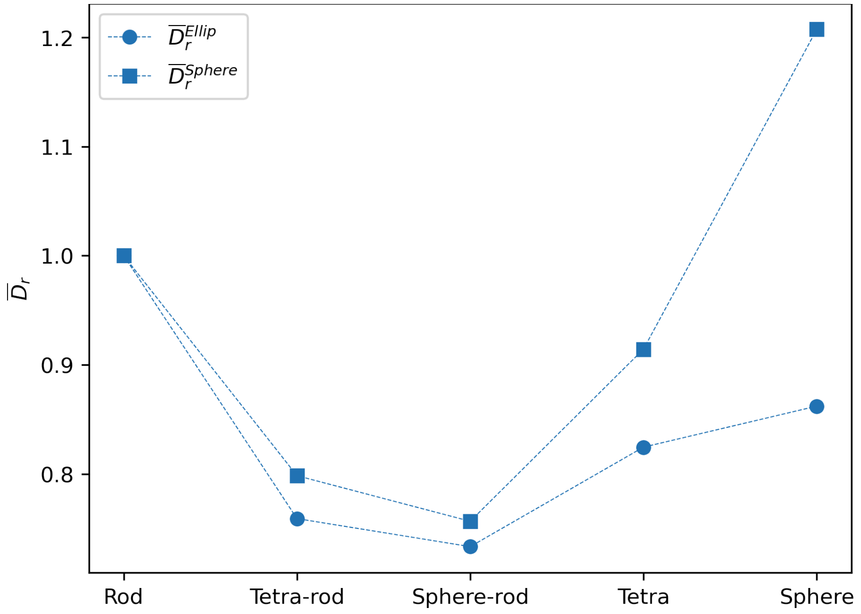


Figure 15: Dimensionless rotational diffusivity for different type of spikes. Diffusivities are calculated using the ratio between the type of spike and the rod spike. Except for the sphere spike in a sphere envelope, all diffusivities are lower than the rod spikes.

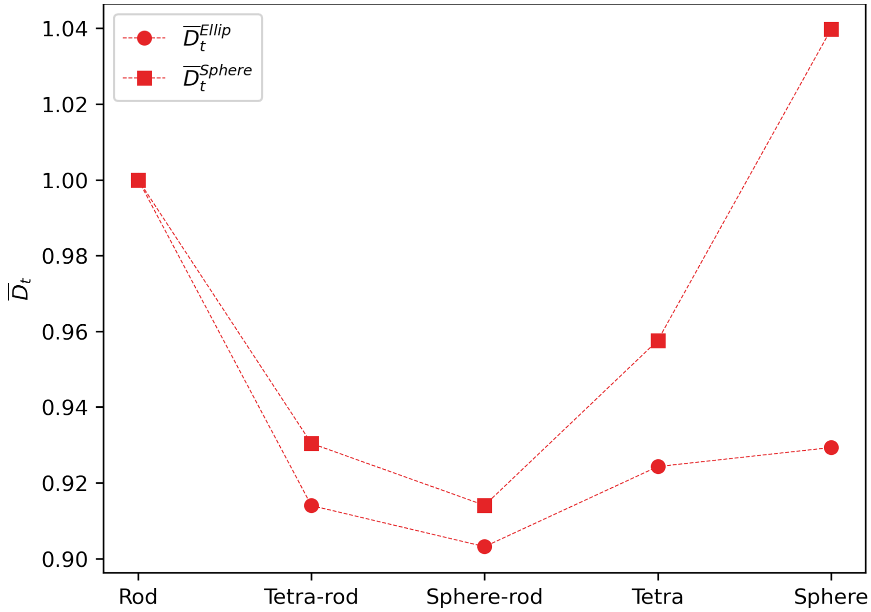


Figure 16: Dimensionless translational diffusivity for different type of spikes. Diffusivities are calculated using the ratio between the type of spike and the rod spike. Except for the sphere spike in a sphere envelope, all diffusivities are lower than the rod spikes.

Table 15: Parameters for simulation for different types of virus.

| **Name** | **Symbol** |
|---|---|
| Biding constant affinity | $K_D$ |
| Translational mobility for ellipsoid | $M_t\|_{\text{ellip}}$ |
| Rotational mobility for ellipsoid | $M_r\|_{\text{ellip}}$ |
| Translational mobility for tetra spike | $M_t\|_{\text{tetra}}$ |
| Rotational mobility for tetra spike | $M_r\|_{\text{tetra}}$ |
| Cell receptor | $\epsilon_{\text{bind}}$ |
| Coefficient of determination | $R^2$ |
| Damkohler number | $D_a$ |
| Dimensioneless rotational diffusivity | $\bar{D}_r$ |
| Dimensioneless translational diffusivity | $\bar{D}_t$ |
| Distance between points of the structure | $ro$ |
| Energy values in molecular-scale systems | $k_B T$ |
| Envelope | $\boldsymbol{E}$ |
| Envelope translational diffusivity | $D_t\|_{\text{env}}$ |
| Envelope rotational diffusivity | $D_r\|_{\text{env}}$ |
| Equivalent radius | $R_e$ |
| Geometrical constraint avidity | $\bar{\Gamma}$ |
| Height of tetra spikes | $H_{\text{tetra}}$ |
| Infectivity parameter | I |
| Lenght of the spike | $l_s/R$ |
| Maximum number of spikes | $N_s^\infty$ |
| Number of spikes | $N_s$ |
| Radius | $R$ |
| Radius of the spike. Pamplomer radius | $p_s/R$ |
| Reaction time | $t_I$ |
| Reaction time biding | $t_b$ |
| Reaction time diffusional | $t_d$ |
| Rotational Diffusivity | $D_r$ |
| Rotational virion diffusivity | $D_r^{\text{virion}}$ |
| Spike proteins | $\boldsymbol{S}$ |
| Temperature | T |
| Temperature of the blood | $T_b$ |
| Theoretical rotational diffusivity | $D_r^o$ |
| Theoretical translational diffusivity | $D_t^o$ |
| Theoretical radius of the envelope | $R_t^o$ |
| Time of rotational | $T_b$ |
| Temperature of the blood | $T_b$ |

| | |
|---|---|
| Traslational Diffusivity | $D_t$ |
| Traslational virion diffusivity | $D_t^{\mathrm{virion}}$ |
| Viscosity | $\eta$ |
| Viscosity of the blood | $\eta_b$ |
| Volume of ellipsoid | $V_{\mathrm{ellip}}$ |
